# Multi-omics and spatial analysis of microgravity-grown glioblastoma organoids reveals superior modeling of advanced disease after long-term spaceflight

**DOI:** 10.64898/2026.03.06.710192

**Authors:** Alice Burchett Darantiere, Maksym Zarodniuk, Shelby Giza, Jason Rexroat, Paul Kuehl, Twyman Clements, Keerthiveena Balraj, Julian Najera, Rohit Bhargava, Meenal Datta

## Abstract

Glioblastoma (GBM) is an incurable brain cancer characterized by its highly immunosuppressive tumor microenvironment and aggressive malignant features that resist treatment. To overcome limitations of Earth-based models (sedimentation and disaggregation) and leverage the unique biological effects of space (accelerated disease progression and immune dysregulation), we developed a panel of GBM-myeloid organoids for extended culture on the International Space Station. After 40 days, the spaceflight-grown organoids had more uniform and reproducible morphology compared to identical ground controls. Organoids containing GBM cells + monocytes had increased expression of chronic innate inflammation, adaptive immune activation, and tissue and vascular remodeling-associated genes. There was an increase in organization of gene expression patterns, with mesenchymal-related genes enriched in the core and inflammation-related genes enriched at the periphery, mimicking GBM tumor architecture. Secretomics confirmed the generation of more immunosuppressive organoids, with enrichment of proteins associated with more aggressive disease, including CXCL12 and LOX-1. GBM co-culture organoids thus had enhanced transcriptomic, proteomic, and architectural features when grown in microgravity that are associated with worse patient outcomes from retrospective data. Infrared laser scanning microscopy confirmed spatial chemical gradients for DNA, protein, and lipid species in both space- and terrestrially-grown organoids. In summary, we present not only a novel and superior model of glioblastoma for more relevant basic, mechanistic, and translational research, but also demonstrate methods to acquire high-quality and diverse data from organoids compatible with the unique experimental constraints of biological research in space to help establish a working model for orbital oncology.

## INTRODUCTION

Microgravity presents an intriguing paradox in the context of human health. As a designated hazard of spaceflight, it alters human physiology and poses a threat to crew health and performance. Conversely, microgravity is used as an advantageous environment for biomedical research to the benefit of humans on Earth, enabling modeling of health and disease in a manner unachievable under Earth’s gravity [1]. In particular, the accelerated development of human-recapitulatory disease states observed in human tissue constructs grown in microgravity – in the absence of gravitationally-induced cell and tissue culture artifacts – renders it a valuable opportunity for time-sensitive pathologies such as glioblastoma (GBM). Despite recent advances in GBM therapies, overall survival remains less than two years [2]. More effective therapeutic strategies are urgently needed, and new approach methodologies (NAMs), such as cancer organoids, offer an avenue towards improved drug discovery in GBM. While organoid modeling techniques for understanding and treating GBM are becoming more widely adopted, they are inherently limited in their ability to recapitulate GBM features and are subject to settling, sedimentation, and disaggregation under Earth’s gravity – as are any human 3-D tissue/disease models. Thus, microgravity-grown organoids are emerging as a promising tool for disease research and therapeutic development [3].

Recent studies have leveraged the microgravity conditions of the International Space Station (ISS) to develop enhanced neural organoids, finding that the organoids in microgravity had decreased markers of proliferation and increased markers of maturation and neuroinflammation, prompting further research to determine if microgravity also promotes accelerated modeling of neurodegenerative diseases [4], [5], [6], [7]. Others have observed hyperplasia in a subset of neural stem cells in 2D culture after 40 days in space, with abnormal cell division resembling that of cancer [8]. Markers of accelerated aging-like phenotypes, such as potential telomere shortening, are observed in blood cells and hematopoietic stem and progenitor cells, both *in vitro* and in astronauts [9], [10], [11]. Microgravity can also cause increased activity of cancer drivers such as ADAR1, highlighting space as a valuable platform to test a novel targeted inhibitor, now in a phase 1 clinical trial [9], [10], [12]. *In vivo* studies on the ISS demonstrate signs of neurodegenerative disease and neuroinflammation in the brain tissues of mice after several weeks in space [13]. After only a few minutes of microgravity exposure on a parabolic flight, patient-derived GBM cells acquired a more aggressive, migratory phenotype with increased growth both *in vitro* and *in vivo* observed after return to ground [14]. However, studies published to date on GBM cells have used either short-duration suborbital flight (16 minutes between launch and landing) or simulated microgravity [3], [14].

When considering GBM organoid design, the immune compartment is of major relevance. Roughly half of the GBM tumor microenvironment is comprised of tumor-associated myeloid cells, including monocytes, macrophages, and microglia, which thwart effective anti-tumor adaptive immune responses and promote treatment resistance, disease progression, ultimately contributing to poor patient outcomes [15], [16]. Macrophages also represent opportunities, as reprogramming or re-engineering them can contribute to therapeutic efficacy [17], [18], [19]. Astronauts also experience immune dysregulation, particularly after prolonged microgravity exposure, characterized by increased innate immune presence and activity and decreased adaptive immune efficacy [20], [21]. Among peripheral blood cell types, monocytes may be the most profoundly impacted by spaceflight [20]. Macrophages exposed to microgravity have impaired differentiation, quantity, and functional polarization compared to 1*g* controls across a range of *in vivo, in vitro,* real, and simulated microgravity experiments [22], [23], [24]. After several days in microgravity, macrophages exhibit a disrupted physical structure, with less actin, more tubulin, and impaired cytoskeletal organization [25]. Concurrent downregulation of CD18, CD36, and MHC-II indicates reduced capacity for pathogen recognition and T cell activation [25]. The macrophage response to microgravity may be mediated by the RAS/ERK/NFκB pathway, which is altered in macrophages in microgravity [26], [27]. This innate immune dysregulation and suppression of effective adaptive immune responses observed in microgravity mimics many of the immunosuppressive characteristics of the GBM tumor microenvironment. Together, these observations suggest that microgravity may be advantageous for GBM modeling due to the simultaneous acceleration of aging-like phenotypes and disease progression with innate immune dysregulation.

Here, we report a first-of-its-kind study on the effects of long-term spaceflight on GBM-immune organoids (**Figure 1**). We successfully optimized methods for the static maintenance and downstream multi-omics analysis of organoids in Space Tango’s autonomous, High Sample Capacity CubeLab platform, which launched on SpaceX’s 30^th^ commercial resupply service flight (SpX-30) to the ISS. After 40 days of growth in space (36 days on ISS plus transit time), an array of GBM and GBM+myeloid organoids was returned to Earth and analyzed for their morphological, transcriptional, and proteomic response to microgravity. GBM organoids grown in microgravity were more uniformly shaped and compact, with less disaggregation compared to identical and simultaneous control organoids (ground controls) maintained at the Kennedy Space Center (KSC). Microgravity induced a distinct transcriptional response depending on the cellular content of the organoids, with GBM alone organoids downregulating mesenchymal-like signatures, and GBM+monocyte organoids upregulating genes associated with chronic innate inflammation, adaptive immune activation, and tissue and vascular remodeling. Spatial transcriptomics revealed that microgravity induces an increase in spatial autocorrelation and radial gradients of gene expression patterns, with mesenchymal-related genes generally enriched in the core and inflammation-related genes enriched at the periphery, mimicking human GBM tumor architecture [28]. Secretomics confirmed the generation of more aggressive organoids, with enrichment of proteins associated with pathological features of GBM, including CXCL12, LOX-1, IL13, and IL17A. Finally, using infrared spectroscopic imaging and associated chemometric analysis, we compared organoids for whole-chemical signatures, confirming spatial patterning of nucleic acids, proteins, and lipids in space- and terrestrially-grown organoids. Together, these results demonstrate a novel model of advanced, immunosuppressive GBM, laying the groundwork for future GBM drug screening and broader disease modeling in space.

**Figure 1.**
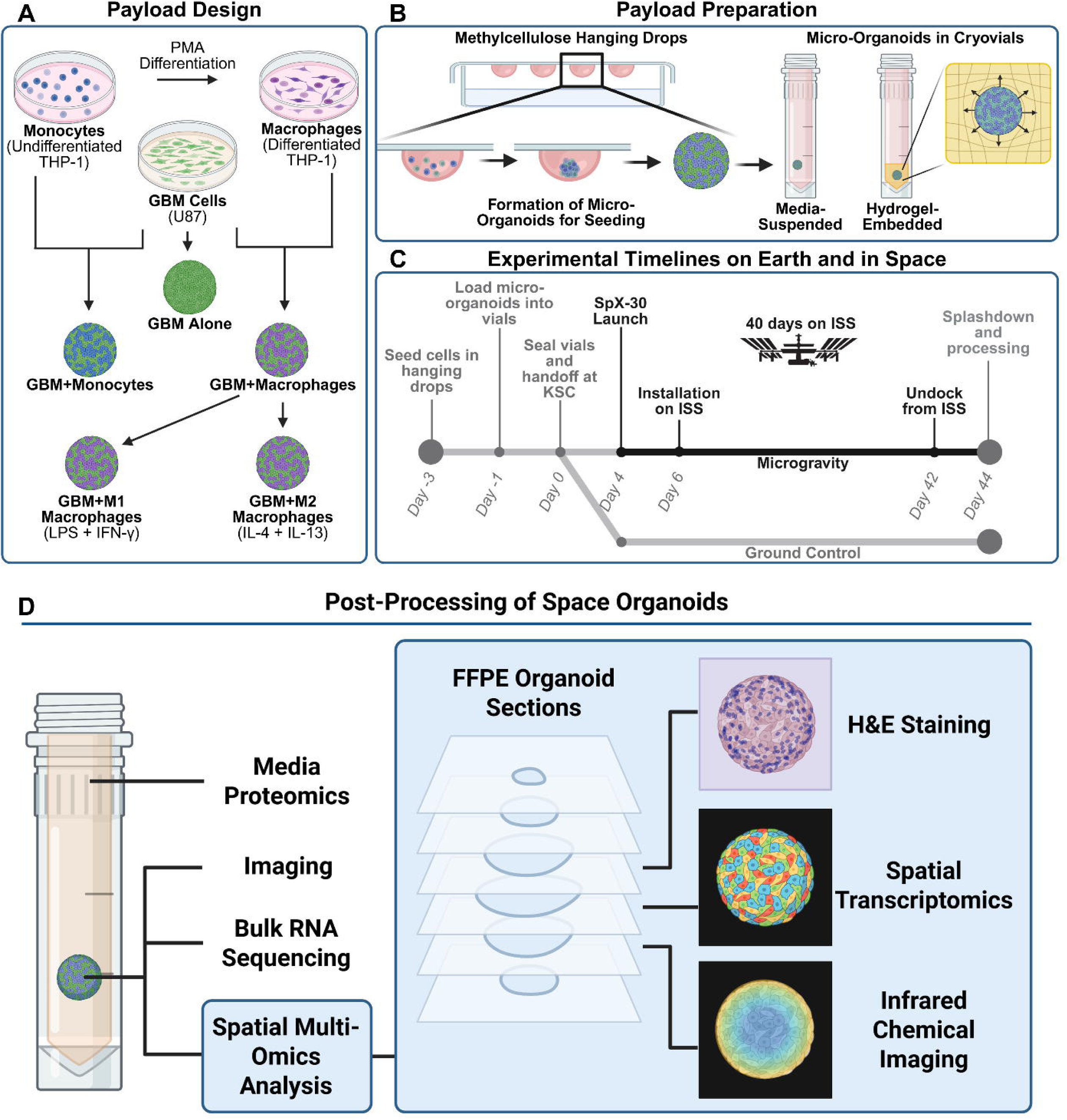
Overview of GBM organoid conditions and methods of payload preparation and analysis. (**A**) Organoids were created with three distinct cellular compositions – GBM cells alone, GBM+monocytes, and GBM+macrophages in 1:1 mixtures. GBM+macrophage organoids were further treated with cytokines to polarize the macrophages into an M1-like or M2-like phenotype. (**B**) Overview of the hanging drop method of organoid formation (left) and schematic of organoids either free-floating or embedded in agarose in cryovials (right). (**C**) Timeline of the spaceflight experiment from organoid formation to endpoint processing. (**D**) Overview of post-flight multi-omics analysis methods of the organoids.

## MATERIALS AND METHODS

### Cell Culture

Human GBM U87 cells and human monocyte THP-1 cells were purchased from ATCC (#HTB-14 and #TIB-202). All cells used in spaceflight experiments were between the 3^rd^ and 5^th^ passages. Cell lines were verified to be free of human pathogens by an external pathogen testing service provider (IMPACT IV PCR Profile plus LCMV, IDEXX Bioanalytics). U87 and THP-1 cells were grown in complete culture medium composed of DMEM (Corning, 10-013-CV) supplemented with 10% fetal bovine serum (FBS, Gibco, 26140079) and 1% penicillin/streptomycin (Corning, 30-001-CI) at 37°C, with 20% O_2_ and 5% CO_2_.

To differentiate THP-1 monocytes into adherent THP-1 macrophages, THP-1 cells were cultured in complete medium supplemented with 10 ng/ml phorbol 12-myristate 13-acetate (PMA, Sigma-Aldrich, P8139-1MG) for 48 hours. This was followed by a 24-hour rest period in regular complete culture medium. Successful differentiation was verified by cells becoming adherent rather than growing in suspension.

### Organoid formation

Cells were seeded in hanging drops to form monoculture or co-culture organoids. Cells were suspended in complete culture medium with 2.4 mg/ml methylcellulose (4000 cPs viscosity, Thermo Scientific Chemicals, 036718.36). The cells were seeded at a range of densities from 50 to 250 cells per 25 µl droplet. To form the hanging drops, 25 µl of the cell-methylcellulose suspension was pipetted onto the inside surface of the lids of 60 mm sterile petri dishes. Sterile water was added to the bottom of the dishes to prevent droplet drying, and the lids with droplets were gently placed on top. These were cultured under standard incubation conditions for 48 hours to allow cell aggregation. Monoculture organoids were composed of U87 cells alone, and co-culture organoids were composed of a 1:1 mixture of U87 cells and THP-1 macrophages (differentiated) or monocytes (undifferentiated), with 250 total cells per drop.

### Fluorescence and brightfield imaging of whole organoids

For visualizing organoid formation in hanging drops, the lid of the petri dish with the droplets was gently inverted and placed on the microscope stage. Images were taken using a brightfield cell culture microscope (Leica DMi 1).

For staining for incorporation of immune and GBM cells, THP-1 monocytes or macrophages were first stained using a live cell membrane stain (CellBrite Blue, Biotium, 30024) before incorporation into hanging drops. After 48 hours, the organoids were stained with Calcein-AM and imaged (Nikon AXR), with maximum intensity Z projections used for representative images.

For viability staining, organoids were transferred to a 96-well plate and stained with Calcein-AM (1:500, BioLegend, 425201) and propidium iodide (2 µg/ml, Biotium, 40016) for 30 minutes at 37°C. Z-stack images of each entire organoid were taken using a confocal microscope (Nikon AXR), and a maximum intensity projection was generated for visualization of representative organoids.

### Loading organoids into cryovials

After 48 hours, organoids were transferred from the hanging drops into cryovials (Thermo Scientific, 374088), to be grown either freely suspended in medium or embedded in agarose. For the free-floating condition, one organoid droplet was transferred to each cryovial, containing 1 ml of culture medium. Culture medium was either DMEM + 10% FBS, DMEM + 1% FBS, or complete NeuroCult medium. Complete NeuroCult medium (NC) is composed of NeuroCult NS-A proliferation kit (STEMCELL Technologies, 05751) supplemented with 20 ng/ml EGF (STEMCELL Technologies, 78006.1), 20 ng/ml FGF (STEMCELL Technologies, 78003.1), and 0.0002% heparin (STEMCELL Technologies, 07980). For the M1 polarized condition, co-culture U87 + THP-1 macrophage organoids were grown in vials with culture medium supplemented with 20 ng/ml of IFN-γ (BioLegend, 570202) and 100 ng/ml lipopolysaccharide (LPS, Santa Cruz Biotechnology, sc-3535). For the M2-like condition, complete medium was supplemented with 20 ng/ml IL-4 (BioLegend, 574002) and 20 ng/ml IL-13 (BioLegend, 571102).

For the agarose-embedded condition, a 4% agarose solution was made with low-gelling temperature agarose (Sigma-Aldrich, A0701-25G) in complete Neuro-Cult medium. The agarose was dissolved by immersion in boiling water, with short bursts in a microwave as needed. The solution was then diluted to a 1% final concentration in warm complete NeuroCult medium and held at 48°C before use. 100 µl of molten agarose was added to each cryovial. One organoid was then pipetted into each vial into the agarose. After gelation, 900 µl of medium (as described above) was added to each vial for a total volume of 1 ml. A summary of the organoid conditions is provided in **Supplementary Table S1**.

To equilibrate the media with the standard 20% O_2_ and 5% CO_2_ gas of the cell culture incubator, the vials were left partially unsealed by turning the lids one-half turn (180°) counterclockwise from the tightened position. After overnight equilibration, the vials were sealed by completely tightening the lids.

### Organoid maintenance and transportation to ISS

The sealed cryovials were transported by vehicle from the University of Notre Dame to the Kennedy Space Center (KSC) in a portable powered incubator (CellBox Solutions) and handed over to Space Tango to be included in the SpaceX Commercial Resupply Services mission CRS-30 in March of 2024 (SpX-30). The samples were integrated into the High Sample Capacity CubeLab and loaded into the SpaceX Falcon-9 rocket three days before liftoff. The exterior surface of each vial was wiped with 70% isopropyl alcohol before loading into the CubeLab. All experimental conditions were duplicated, with one set loaded into a CubeLab unit scheduled for launch, and the other maintained in an incubator at KSC for ground controls at Earth’s gravity. Prior to NASA handover, the CubeLab was purged with 5% CO_2_ (balance breathing air). The CubeLab was loaded into Space Tango’s Powered Ascent Utility Locker (PAUL), which provided power and data to the CubeLab during all phases of the mission, maintaining 37°C temperature for organoid viability. Two days after launch, the Dragon capsule docked to the ISS, and the Cubelab was installed on the ISS. After 36 days on the ISS, the CubeLab was placed in the Dragon cargo capsule, which undocked for return to Earth, splashing down two days later in the Atlantic Ocean off the coast of Florida. With the exception of one brief anomaly, the CubeLab system maintained the sample temperature at 37°C and provided passive CO_2_ gas from the time of handoff to the time of sample retrieval.

Within five hours after Dragon splashdown, the live samples were retrieved from the ISS CubeLab (de-integrated at KSC’s Space Station Processing Facility [SSPF]) and KSC ground control incubator and transported back to the University of Notre Dame by flight for processing. The samples were maintained near 37°C in an insulated box during transit with chemical heating packs (ThermaCare) and were placed in a standard cell culture incubator on their arrival. All samples were processed and preserved within 24 hours of splashdown.

### Radiation monitoring

Radiation data was collected from the Radiation Environment Monitor (REM) nearest to the location of the CubeLab payload. The galactic cosmic radiation (GCR) and Southern Atlantic Anomaly (SAA) detected dose was recorded daily (mGy/day) from the day the payload was launched to the day the payload was retrieved (**Supplementary Table S2**). This data was provided by the NASA Space Radiation Analysis Group. GCR and SAA readings were combined to yield the average daily radiation and cumulative mission radiation exposure.

### Collection of conditioned media for proteomics analysis

Color photographs were first taken of representative vials of each condition to document media acidification (Pixel 7a). Vials with free-floating organoids were centrifuged at 200xg for 5 minutes, and the media was carefully removed and stored at −80°C. For vials with agarose-embedded organoids, the media was carefully removed to avoid disturbing the agarose gel. For the media-only controls, leftover media after organoid loading into vials was stored at 4°C, then briefly frozen at −80°C, and prepared in the same way as the conditioned media. Media samples were thawed on ice, inverted to mix, and centrifuged at 3,000 x g for 5 minutes to remove debris. The supernatant (∼80% of the original sample) was collected, inverted again to mix, and 150 µl was removed for proteomics analysis. Samples were frozen again and shipped on dry ice to Azenta Life Sciences for analysis via the Olink^®^ Proximity Extension™ Assay Target 48 Cytokine panel.

### Fixation and paraffin embedding of whole organoids

After the media was removed from agarose-embedded organoids, the remaining volume of the cryovial was filled with 4% paraformaldehyde in PBS (Thermo Scientific Chemicals, J19943.K2), and the organoids were fixed overnight at 4°C. After several PBS washes, the cryovials were filled with 70% ethanol and stored at 4°C for later analysis.

Representative organoids next underwent FFPE processing and sectioning. First, individual organoids (agarose-embedded condition) were dissected out of the agarose gel using a razor blade. The organoids were placed into the bottom of cone-shaped tissue molds, and all excess liquid was removed. A warm solution of 2% low gelling-temperature agarose plus 2% gelatin (Sigma Aldrich, 2500-100G) in PBS was added on top and manually mixed to ensure that the organoid was positioned at the tip of the conical tube but still surrounded by agarose. After the gel had solidified, the samples were stored in 70% ethanol before being processed and embedded in paraffin (FFPE) blocks.

### Organoid morphology quantification

For quantitative assessment of endpoint morphology, one organoid from each agarose-embedded condition was dissected out of the agarose gel after fixation and imaged using a standard brightfield microscope (Leica DMi 1). Images with artifacts preventing accurate organoid segmentation were excluded. Automated segmentation of the organoid images was performed using CellProfiler, which identified the organoid in each image and reported its solidity and form factor. Solidity is one metric of organoid irregularity and is defined as the area of the object divided by the area of the convex polygon that would be needed to contain the whole object (Area ÷ Convex Hull Area), such that a perfect circle has a solidity value of 1, and irregular shapes have solidity values less than 1. Form factor is defined as 4×π×Area÷Perimeter^2^ and is a metric of the smoothness of an object’s boundary, with a perfect circle corresponding to a form factor of 1.

### Xenium Spatial Transcriptomics Assay

FFPE blocks containing the organoids were stored at 4°C prior to sectioning. Four conditions from the KSC ground controls and four conditions from microgravity were used for spatial transcriptomics: U87 alone (GBM alone), U87+differentiated THP1 macrophages (GBM+MΦ), U87+undifferentiated THP1 monocytes (GBM+mono), and U87+M2-like polarized THP1 macrophages (GBM+M2). One 6 µm-thick section from each of the eight organoids was included in the capture area of a single Xenium slide (10x Genomics, 1000465). The sections were adhered to the slide sequentially after rehydration in an RNase-free water bath.

Samples were processed using the 10x Genomics Xenium Prime 5K Human Pan Tissue and Pathways Assay Kit (PN 1000671; 10x Genomics, Inc.) in combination with the Xenium Prime 5K Human Pan Tissue & Pathway Panel (PN 1000724; 10x Genomics, Inc.). Tissue preparation, probe hybridization, signal amplification, and optional cell segmentation staining were performed according to the manufacturer’s protocols as described in the Xenium Prime In Situ Gene Expression with Optional Cell Segmentation Staining user guide (Doc# CG000760 Rev C) and the Xenium In Situ - FFPE Tissue Preparation Handbook (Doc# CG000578 Rev E).

Imaging and data acquisition were conducted using the 10x Genomics Xenium Analyzer Instrument (Doc# CG000584 Rev J) with Instrument Controller Software v3.3.0.1 and on-instrument analysis software v3.3.0.1. A total of eight organoids were imaged, with each organoid individually selected and analyzed as a single region of interest (ROI) on the slide.

Serial sections of each organoid were stained with hematoxylin and eosin (H&E) to validate sample quality and visualize the necrotic core.

### Bulk RNA extraction and sequencing

After the media was removed from the vials with free-floating organoids, the remaining organoid was left in the original cryovial, flash-frozen in liquid nitrogen, and stored at −80°C. RNA was isolated from each organoid individually. The organoids were thawed slightly on ice and gently flicked to loosen. They were then resuspended in 350 µl of buffer RLT (Qiagen, 79216) and vortexed for 2 minutes at 3,000 rpm. The RNA was isolated using a Qiagen RNeasy micro kit (Qiagen, 74034) and eluted in 12 µl of RNase-free water. RNA quality and quantity were assessed using a TapeStation HS RNA assay, with lower-abundance samples quantified again using the Bioanalyzer RNA Pico Assay.

Stranded Illumina RNA-seq libraries were prepared from 250 pg to 1 ng total RNA using the SMART-Seq Total RNA Pico Input with UMIs kit (ZapR Depletion, Mammalian) (Cat# 634354, Takara Bio USA Inc; User Manual #052424) and the Unique Dual Index Kit (Cat# 634354, Takara Bio USA Inc., User Manual #052424), following the manufacturer’s protocol with the modifications described below. Fragmentation time was selected based on the higher reported eRIN (TapeStation High Sensitivity RNA ScreenTape Assay, Cat# 5067-5579) or RIN (Bioanalyzer RNA Pico Chip Assay, Cat# 5067-1513) (Agilent Technologies, Inc.). For samples with undetermined eRIN/RIN values, fragmentation time was empirically determined by constructing a library with 0 min fragmentation and comparing the resulting fragment profile to a reference library of known eRIN/RIN generated using titrated fragmentation times. The PCR2 cycle number was determined by estimating total RNA concentration using the TapeStation High Sensitivity RNA ScreenTape Assay. For samples with an undetermined total RNA concentration, PCR2 cycles were empirically optimized by preparing a library with 2 µL sample input and titrating PCR2 cycles to achieve a 4 nM to 100 nM final library concentration.

For multiplexing, library molarity was determined for each library using the KAPA Library Quantification Kit for Illumina (Cat# 07960140001) and the TapeStation High Sensitivity D5000 ScreenTape Assay (Cat# 5067-5592). Libraries were sequenced on an Illumina NovaSeq X System using an XLEAP 10B flow cell with 2 × 100 bp paired-end reads, targeting 120M clusters (240M paired reads) for microgravity samples and 60M clusters (120M paired reads) for ground control samples. Real-time base calling and demultiplexing were performed on-instrument using DRAGEN software to generate gzipped FASTQ files (2 × 100 bp). Library preparation was performed by the University of Notre Dame Genomics and Bioinformatics Core Facility (Notre Dame, IN), and sequencing was performed by the Indiana University School of Medicine Center for Medical Genomics (CMG) (Indianapolis, IN).

### Bulk RNA-seq data preprocessing and analysis

UMI-based bulk RNA-seq data were processed using a custom analysis pipeline. Raw paired-end FASTQ files were first processed to extract unique molecular identifiers (UMIs) using UMI-tools (v1.1.6; extract). UMI-processed reads were subsequently trimmed with Trimmomatic (v0.39) to remove the SMART adapter and technical sequences.

Trimmed reads were aligned to the GRCh38.p14 genome assembly (GCF_000001405.40) using STAR (v2.7.2), generating coordinate-sorted BAM files. BAM files were indexed with samtools (v1.20) to facilitate downstream processing and quality control. PCR duplicates were removed using UMI-tools (v1.1.6; dedup), which performs UMI-aware read grouping based on genomic alignment position and UMI sequence similarity. Deduplication statistics were collected to assess UMI complexity and duplication rates. Gene-level expression quantification was performed on deduplicated BAM files using Subread featureCounts (v2.0.8).

Technical replicates were collapsed by summing raw counts. Differential expression analysis was conducted using DESeq2 (v1.46.0) with a two-factor interaction model incorporating cell composition (with vs. without monocytes) and experimental condition (uG, microgravity vs. KSC, Kennedy Space Center ground controls). Differentially expressed genes (DEGs) were defined as those with Benjamini-Hochberg-adjusted p values < 0.05 and were classified as up- or downregulated based on the direction of the DESeq2-estimated log₂ fold change.

Gene set enrichment analysis (GSEA) and over-representation analysis (ORA) were performed using clusterProfiler (v4.14.6). GSEA was conducted using DESeq2 Wald test statistics as the ranking metric unless otherwise specified. ORA was performed separately for up- and downregulated gene sets, with the universe defined as all genes present in the original count matrix. Pathways with ORA p values < 0.05 and q values < 0.2 were considered significantly enriched.

Motif enrichment analysis was performed using HOMER (v5.1; findMotifs) by scanning promoter regions spanning 400 bp upstream to 100 bp downstream of transcription start sites (TSSs). Motif lengths of 8 and 10 base pairs were tested. Genomic locations of the top three ETS-family motifs were identified using the annotatePeaks module in TSS mode.

For survival analysis, GBM RNA-seq data from the Cancer Genome Atlas (TCGA) was deconvolved using BayesPrism with a published single-cell reference (GSE84465) [29]. Outlier genes were removed using BayesPrism’s built-in filtering; the reference was further restricted to protein-coding genes and informative markers. Bulk counts were filtered to retain expressed genes and supplied to new.prism, followed by model fitting with run.prism. Malignant (Glioma) expression profiles inferred by BayesPrism were extracted and variance-stabilized for survival analysis.

We then computed single-sample gene set enrichment scores (ssGSEA) from variance-stabilized expression data using the top 50 genes based on Wald statistic. Scores were mean-centered and scaled, averaged per patient, and stratified into “high” vs “low” groups using the median. Associations with overall survival and progression-free survival were tested using a multivariable Cox proportional hazards model including the ssGSEA signature and covariates (age, sex, MGMT promoter status, and IDH status).

### Xenium in situ transcriptomics data filtering

Xenium spatial transcriptomics data from eight organoids were integrated into a single SpatialData object (spatialdata 0.4.0). Cells were filtered based on transcriptome complexity, retaining only those for which fewer than 55% of total transcript counts were contributed by the 50 most highly expressed genes.

### Consensus NMF (cNMF) analysis

cNMF was performed using Python implementation of cnmf (1.7.0) on the merged and filtered Xenium gene expression matrix. To identify an appropriate number of gene expression programs (GEPs), we first conducted a parameter sweep over the number of components K, evaluating values from K = 2 to K = 10 with 50 NMF iterations per K. Based on the sweep, we selected K = 4 for downstream analyses and reran cNMF with increased consensus stability by performing 500 iterations.

### GEP signature scoring

GEP activity was quantified using signature scores, as normalized cNMF usage reflects relative program composition and may mask absolute transcriptional upregulation when programs co-vary. For each GEP, we defined a “GEP signature” as the top 50 genes (ranked by GEP loading) and calculated a per-cell “GEP signature score” based on the relative expression of genes in that signature compared to background genes matched for expression level, as implemented in sc.tl.score_genes.

### Assignment of cells to GEPs

To assign individual cells to one or more GEPs, we used a permutation-based framework built on the GEP signature scores, as described previously [30]. First, we constructed an empirical null distribution for each GEP score by permuting the expression matrix 100 times. In each permutation, gene expression values were randomly shuffled across cells, preserving each gene’s overall distribution while disrupting covariance structure. For every permuted dataset, we recomputed GEP signature scores using the top 50 genes per GEP (as above) and aggregated the resulting null score distributions.

For each observed cell-GEP pair, we then calculated a one-sided empirical p-value by comparing the observed GEP score to the corresponding permutation-derived null distribution (assuming approximate normality of the null, using the null mean and standard deviation). P-values were adjusted for multiple testing using the Benjamini-Hochberg false discovery rate (FDR) procedure. Cells were assigned to a GEP if their adjusted p-value for that GEP was < 0.05.

### Gene set enrichment analysis

GSEA was performed using clusterProfiler (v4.14.6). Enrichment was evaluated using GEP scores (i.e., NMF weights).

### Quadrant plots

Quadrant plots were generated as described previously [31]. Briefly, quadrant coordinates were derived from normalized NMF usage by collapsing the four GEP usage values into two contrast axes and then rotating the space for visualization. Specifically, for each cell, we computed two difference scores: X = GEP1 - GEP2 and Y = GEP3 - GEP4. The resulting (X, Y) coordinates were then rotated by 45° clockwise using a rotation matrix.

### Deconvolution of GEPs in bulk RNA-seq data

Bulk RNA-seq GEP deconvolution was performed by ssGSEA using previously defined GEP gene signatures. Raw counts were processed in DESeq2 and variance-stabilized using the VST transformation. The resulting normalized expression matrix was analyzed with GSVA (v2.0.7) in ssGSEA mode to compute enrichment scores for each sample and GEP gene set, yielding a sample-by-GEP score matrix. Scores were z-scored across samples within each GEP to enable comparisons across conditions.

### Radial analysis of organoids

Radial spatial analysis was performed to quantify cell-state organization as a function of distance from the organoid boundary. For each sample, cell boundary polygons derived from Xenium segmentation were retrieved as spatial geometries. To define a smooth outer organoid contour, all cell polygons were first dilated by 10 µm and merged into a single geometry using a unary union operation. The largest resulting polygon was selected to remove spurious fragments, then further smoothed by sequential dilation and erosion (±20 µm). This final polygon was taken as the organoid outer boundary.

Cell centroids were used to compute the shortest Euclidean distance from each cell to the organoid boundary. Only cells fully contained within the organoid boundary were considered. Distances were normalized by the maximum interior distance for each organoid, yielding a normalized radial coordinate ranging from 0 at the boundary to 1 at the organoid core.

Cells were discretized into three radial layers—periphery, middle, and core—using equal-width bins of the normalized radial coordinate [0, 1/3), [1/3, 2/3), [2/3, 1]. To estimate the total area of each radial layer, cell polygons within each layer were merged after smoothing (±20 µm), and the resulting areas were computed and summarized per sample.

### Generalized additive model analysis of radial GEP gradients

To model non-linear radial trends in GEP activity, we fit generalized additive models (GAMs) to per-cell GEP signature scores as a function of normalized radial coordinate. For each GEP (GEP1-GEP4), we fit a Gaussian GAM of the form: GEP score ∼ s(radial coordinate, by = group) + group, where group is a factor representing organoid condition (i.e. microgravity-grown U87 GBM organoids with monocytes) using mgcv::gam, allowing the smooth radial effect to vary by condition while accounting for baseline group differences. We summarized fitted radial patterns using absolute fitted curves generated by predicting from each model over an evenly spaced grid of radial positions within the central 1st-99th percentile of observed values (to avoid the effect of outliers), with 95% confidence intervals computed as ±1.96*SE.

### Quantum cascade laser (QCL)-based discrete frequency infrared (DF-IR) chemical imaging using a laser scanning microscope

The FFPE organoid samples were sectioned onto an infrared (IR) reflective MirrIR low-e slide (Kevley Technologies) at 6 µm thickness. Prior to imaging, these slides were deparaffinized by soaking in hexane (>98.5%, Fisher Chemical) for 24 hours. We used a custom point-scanning confocal DF-IR microscope that uses a QCL, a thermoelectric cooled mercury cadmium telluride (MCT) point detector, and a 0.71 N.A. refractive objective. Multispectral imaging on the DF-IR system was performed with 2 coadditions and 2 µm pixel size. We emphasize the fingerprint regions (900-1800 cm^−1^), which contain the biologically relevant vibrations such as DNA, protein, and collagens. IR images were acquired on the DF-IR system at 1080, 1240, 1335, 1460, 1540, 1630 and 1658 cm^−1^. The bands between 1600 and 1700 cm⁻¹ fall within Amide I (C=O stretching) and reflect protein secondary structure. The 1540 cm⁻¹ band is typically assigned to Amide II (C-N stretching coupled with N-H bending) region. The 1460 cm⁻¹ band corresponds to lipid/protein content (CH_2_/CH_2_ bending), and the 1335 cm⁻¹ is associated with ECM. The 1240 cm⁻¹ band is often linked to nucleic acids and phospholipids (asymmetric PO₂⁻ stretching), and 1080 cm⁻¹ band is commonly associated with nucleic acid-containing biomolecules (symmetric PO₂⁻ stretching).

The collected DF-IR data were preprocessed to baseline correct for scattering and normalized to control for thickness variations across samples. A binary organoid mask was derived from the reference band and organoids were segmented into core-middle-periphery regions. These radial zones were derived by computing the Euclidean distance to the mask boundary of each pixel. A threshold is applied to create an outer band (periphery), an intermediate ring (middle), and an inner (core) region.

### Statistical Analysis

All statistics were performed either in GraphPad Prism (v11.0.0), R, or Python. The statistical tests are indicated in the figure legends, and all t tests were two-tailed. For morphological analysis, comparisons were made with four ground control images and three microgravity images. For the bulk sequencing data, there were two GBM microgravity samples, four GBM microgravity samples, two microgravity GBM+monocytes samples, and four GBM+monocytes ground control samples. For spatial transcriptomics and DF-IR analysis, there was one organoid for each condition.

## RESULTS

### GBM and myeloid cells successfully form organoids with long-term viability in static cultures

We first developed GBM-myeloid organoids for long-term, static culture experiments compatible with Space Tango’s autonomous, passive hardware system, which has a heritage of flight success [32], [4], [33]. These organoids contained U87 human GBM cells either alone or in combination with THP-1 human monocytes or differentiated THP-1 human macrophages (**Figure 1A**). Because of the high-risk nature of this first demonstration of prolonged GBM organoid growth in space, immortalized human cell lines were selected over primary or patient-derived cells for their robustness, predictability (based on prior terrestrial studies and well-established biological features), and likelihood of durability in long-term spaceflight. We used the hanging drop method to generate spheroidal cell aggregates with tunable cell content (micro-organoids), which can be transferred to vials of culture medium, with the optional addition of cytokine stimulation and/or hydrogel (1% agarose w/v mimicking soft tissues, as in our prior work) embedding (**Figure 1B**) [34], [35]. The addition of methylcellulose to the droplet solution increases its viscosity, stabilizing the liquid droplets and promoting the reliable formation of spheroidal cell aggregates [34]. The GBM organoids were optimized for long-term culture studies to accommodate a 30-45-day mission typical of CRS durations (**Figure 1C**) and downstream multi-omics analysis (**Figure 1D**).

After 48 hours in the droplet, GBM cells alone and 1:1 mixtures of GBM cells and monocytes or macrophages coalesced to form spheroidal aggregates that were on average 130 µm in diameter (±11 µm) (**Figure 2A**). Fluorescent labeling of live macrophages and monocytes demonstrates immune cell incorporation throughout the organoids (**Figure 2B**). We next confirmed that these organoids would remain viable in long-term static culture without gas or media exchange, as constrained by the autonomous, static hardware system. After 47 days (simulating a 40-day mission plus several days of transit) in sealed vials, GBM alone, GBM + monocyte, and GBM + macrophage organoids had grown to roughly 600 µm in diameter. The organoids retained viable cells, particularly in the outer rim, while the cells in the organoid cores were primarily dead (**Figure 2C**). Cell culture medium in the sealed vials became more acidic over time as the cells proliferated, observable by a visible shift in color of the phenol red pH indicator (**Supplementary Figure S1A**). This necrotic core, acidic culture medium, and lack of oxygen supply in the static vial system mimics the necrotic, acidic, and hypoxic microenvironment that is typical of GBM tumors [36].

**Figure 2.**
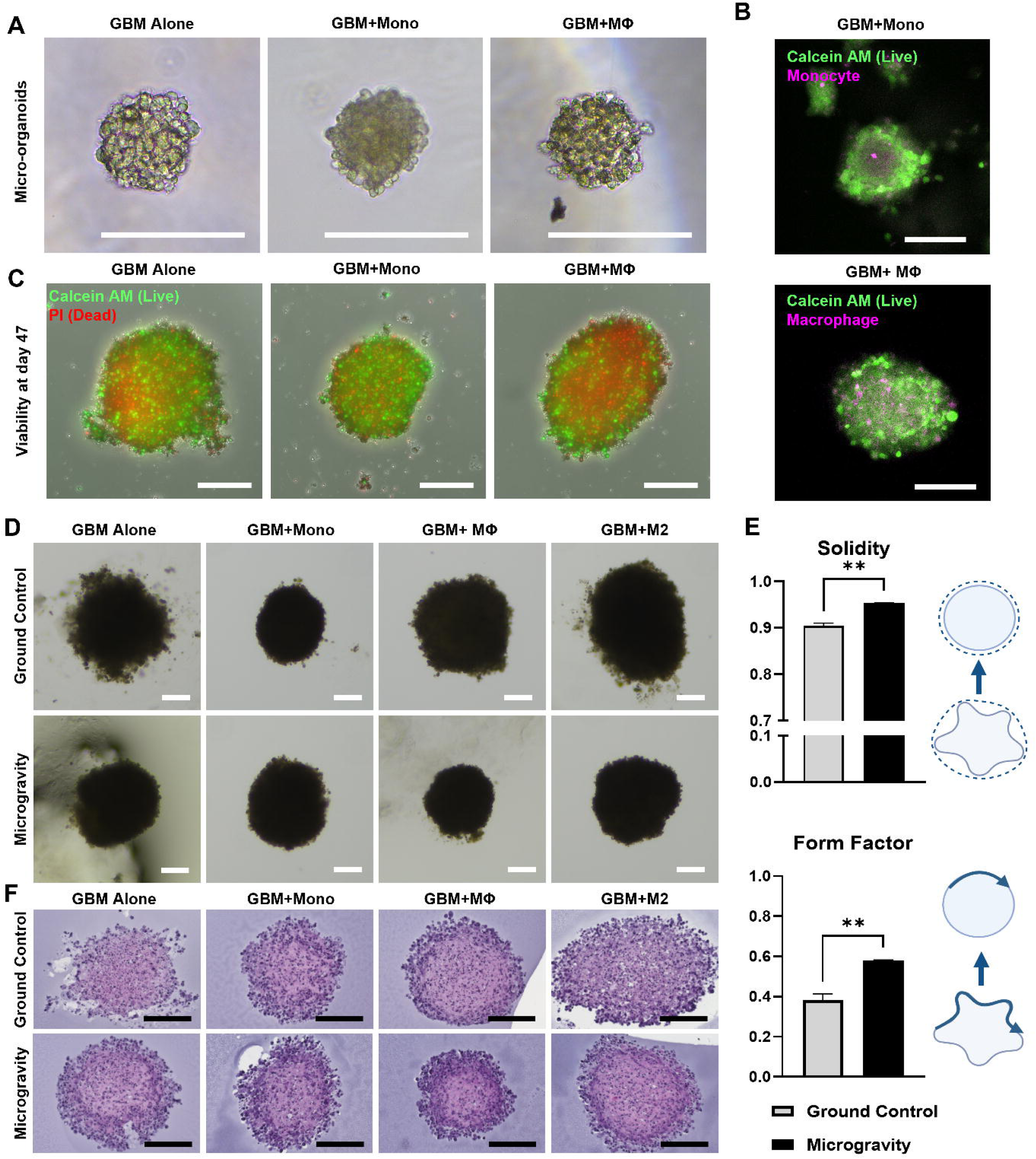
Organoids successfully incorporate immune cells, survive long-term culture, and expand to form more uniform organoids in microgravity than on Earth. (**A**) Phase-contrast images of GBM alone, GBM+monocyte (Mono), and GBM+macrophage (MΦ) organoids after 48 hours in hanging drops. Scale bars represent 200 µm. (**B**) Images showing monocytes and macrophages (magenta) incorporated with GBM cells in live organoids (green). Scale bars represent 200 µm. (**C**) Viability staining showing live cells (green) and dead cells (red) in organoids after 47 days in sealed cryovials. Scale bars represent 200 µm. (**D**) Brightfield images of ground control and post-flight microgravity organoids after fixation and removal from agarose. Scale bars represent 200 µm. (**E**) Quantification of compactness, solidity, and form factor from the brightfield organoid images. Statistical comparisons were made using unpaired t tests (* p < 0.05, ** p < 0.01). Error bars indicate standard error of the mean (SEM). (**F**) Hematoxylin and eosin staining of 6 µm sections of ground control and microgravity organoids. Scale bars represent 200 µm.

The final experimental payload design, supported by Space Tango’s CubeLab system on the SpX-30 mission, included a range of cell types and media conditions (**Figure 1A-B, Supplementary Table S1**). The organoid content variations were U87 cells alone at seeding densities ranging from 50 to 250 initial cells per organoid, U87 cells co-cultured with differentiated THP-1 macrophages at a 1:1 ratio, and U87 cells co-cultured with undifferentiated THP-1 monocytes at a 1:1 ratio. GBM+macrophage organoids were further divided into an unpolarized group grown in standard Neurocult media, an M1-like (anti-tumor, pro-inflammatory) polarized group in Neurocult supplemented with LPS and IFN-γ, and an M2-like (pro-tumor, anti-inflammatory) polarized group supplemented with IL-4 and IL-13. The payload also included U87 and U87+THP-1 organoids in DMEM supplemented with either 10% or 1% FBS.

### Organoids successfully expand in microgravity and have more uniform morphology

Cells were seeded in hanging drops three days prior to the cryovial seal date. After 48 hours, they were transferred to cryovials, either embedded in agarose or freely floating in media, and the vials were cracked open for overnight media equilibration. The organoids were sealed in their vials 4 days before launch, spent 40 days in orbit (36 days on ISS, 4 days in orbital transit), and <24 hours in transit back to our terrestrial lab for post-processing and analysis (**Figure 1C, D**). During this time, an ISS radiation detector that was closest to the CubeLab reported an average daily dose of 0.20 mGy/day of galactic cosmic radiation (GCR), for a cumulative dose of 8.1 mGy (**Supplementary Table S2**). Synchronous ground controls were maintained at KSC in an incubator. All samples were processed within 24 hours of de-integration, immediately following splashdown and retrieval, after having spent a total of 45 days in sealed vials. Upon return, vials with successful organoid formation were identified by an observable change from pink to yellow media color, indicating media acidification due to cell proliferation (**Supplementary Figure S1A**). Vials containing 250 cells as a starting density had more media acidification and cell growth compared to 50 cells and 100 cells (**Supplementary Figure S1B, Supplementary Table S3**); therefore, only the 250-cell samples were analyzed. Agarose-embedded organoids were also more likely to be successful than their free-floating counterparts (**Supplementary Figure S1C, Supplementary Table S3**). This difference was more pronounced in the microgravity condition than in the ground controls. For the microgravity organoids and KSC ground controls, there were almost no successful organoids in the M1-like polarized condition. Organoids grown in DMEM with 10% or 1% also had a low success rate and were excluded from further analysis.

Agarose-embedded organoids grown in microgravity tended to be more cohesive, with fewer visually detached cell aggregates compared to ground controls (**Figure 2D**). Quantification of post-flight brightfield images confirmed statistically significant improvements in organoid morphology (**Figure 2E**). Microgravity-grown organoids had a significant increase in solidity compared to ground controls, indicating they had a more regular, round shape. Microgravity-grown organoids also had a significant increase in form factor, meaning they were smoother and more compact. There was no significant difference in organoid size in microgravity (**Supplementary Figure S2**). Hematoxylin & Eosin staining of the agarose-embedded organoids qualitatively shows intact nuclei in an outer rim, with a necrotic core at the center of the organoid section (**Figure 2F**).

### Microgravity elicits context-dependent transcriptional responses in GBM organoids influenced by monocyte exposure

We performed bulk transcriptomic profiling of media-suspended microgravity-grown GBM organoids with or without THP-1 monocytes using SMART-seq. Differential expression analysis revealed transcriptional changes in response to microgravity in both organoid conditions, with markedly stronger effects in GBM alone organoids and limited overlap in differentially expressed genes (DEGs) between conditions (**Supplementary Figure S3A, Supplementary Table S4**). In both space and on ground, we detected minimal transcriptional differences between organoids cultured with or without THP-1 monocytes (**Supplementary Figure S3A**), suggesting limited persistence of monocytes within organoids after long-term culture (as confirmed later by spatial transcriptomics).

Microgravity-grown organoids derived from GBM cells alone showed reduced expression of genes associated with mesenchymal remodeling pathways, such as extracellular matrix (ECM) organization and angiogenesis, as well as decreased expression of glycolytic and apoptosis-related genes, indicating a shift away from mesenchymal, metabolically glycolytic, and stress-adapted states (**Figures 3A, B**). Instead, they upregulated of genes involved in post-transcriptional regulation, including mRNA processing, nuclear export, and translation.

**Figure 3.**
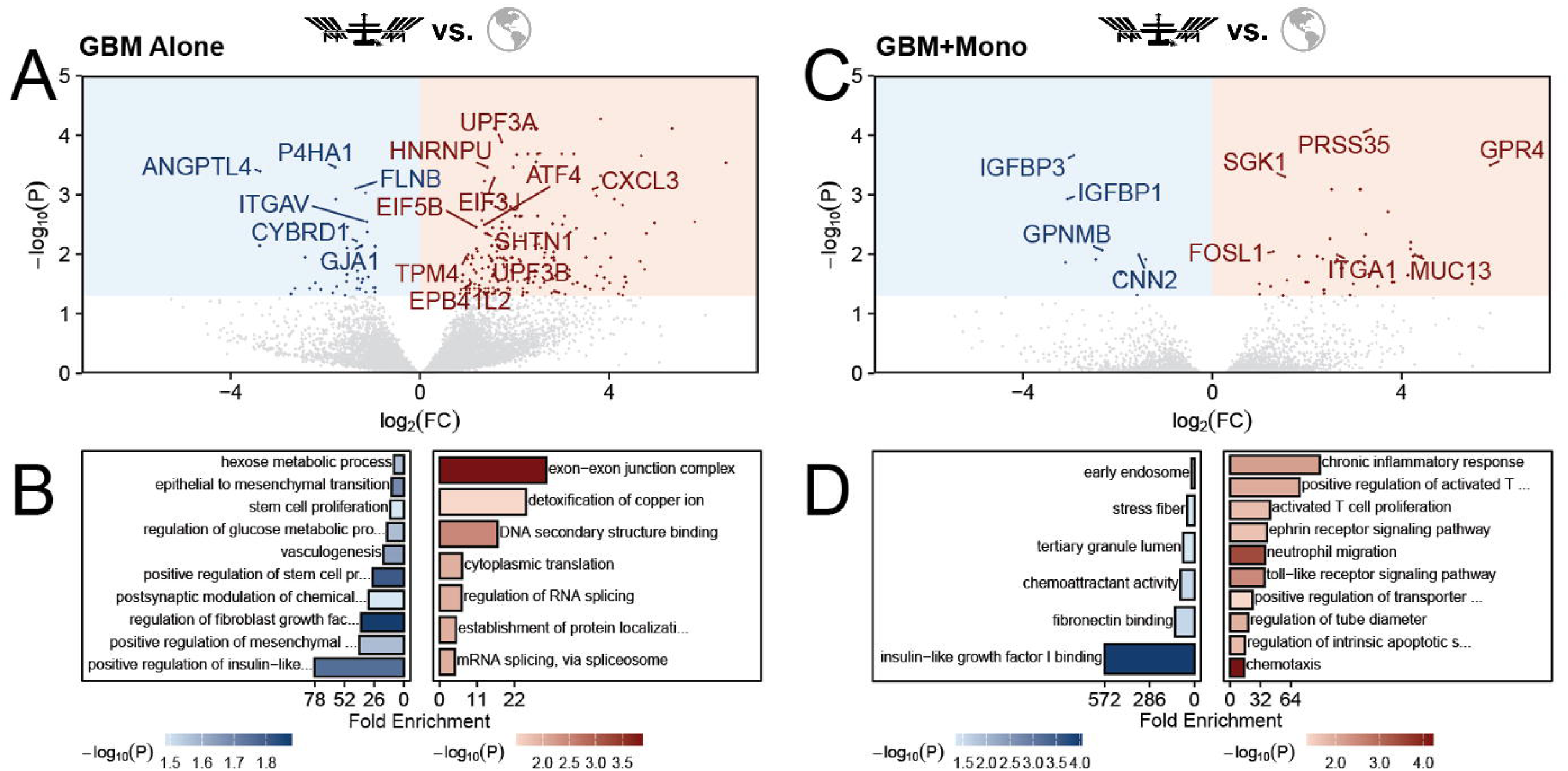
Microgravity elicits context-dependent transcriptional responses in GBM organoids influenced by monocyte exposure. (**A**) Volcano plots show DESeq2 statistics comparing microgravity-grown organoids to Kennedy Space Center (KSC) 1*g* controls for U87 GBM alone (**A**) and GBM co-cultured with THP-1 monocytes (**B**). Each point represents a gene, plotted by log2 fold change (x-axis; microgravity vs control) and -log10 adjusted p-value (y-axis; Benjamini-Hochberg corrected). Genes meeting p < 0.05 are highlighted as significantly upregulated (red) or downregulated (blue); non-significant genes are shown in gray. Shaded regions indicate the significance threshold (P < 0.05; fold-change threshold set to 0), and the top 20 most significant differentially expressed genes in each comparison are labeled. (**C**) Bar plots show Gene Ontology (GO) over-representation analysis (ORA) of genes differentially expressed in microgravity-grown organoids relative to KSC 1*g* controls. Results are shown separately for U87 GBM alone (**C**) and U87 GBM co-cultured with THP-1 monocytes (**D**), with downregulated (blue; left-extending bars) and upregulated (red; right-extending bars) gene sets displayed in separate panels. Bars represent fold enrichment for selected GO terms, and color intensity encodes statistical significance (-log10 adjusted p-value, Benjamini-Hochberg correction). GO terms were simplified to reduce redundancy, and a curated subset of biologically relevant terms is shown for clarity.

In contrast, genes upregulated by microgravity-grown GBM organoids with THP-1 monocytes were associated with chronic innate inflammation, adaptive immune activation, and tissue and vascular remodeling (**Figures 3C, D**). These organoids also exhibited reduced expression of genes involved in insulin-like growth factor I binding, including *IGFBP1* and *IGFBP3* (**Figure 3C**), which are associated with high-grade gliomas and poor clinical outcomes [37], [38]. They also upregulated genes such as *PTGS2* (encoding COX-2) and *S100A8,* reported to be associated with GBM immune dysfunction and advanced disease [39], [40], [41]. *FOSL1* and *ITGA1* were among the up-regulated genes associated with resistance to radiation and temozolomide [42], [43].

Although THP-1 monocytes did not persist in significant numbers after long-term static culture, GBM organoids displayed distinct transcriptional responses depending on initial monocyte incorporation. We therefore hypothesized that monocytes modulate response to microgravity through indirect or cancer cell “preconditioning” effects early during organoid growth. Consistent with this hypothesis, interaction modeling identified several genes with differential microgravity effects (**Supplementary Figures S3B, C**), including *PRSS35* (serine protease 35), *EPHA7* (ephrin receptor A7), and *FAM50A,* which encodes a component of the spliceosome complex C. Together with the limited overlap in DEGs across conditions, these data support a role for monocyte exposure in modulating GBM cell transcriptional responses to microgravity.

### Microgravity-associated transcriptional changes align with GBM meta-program and TEAD/SMAD3 regulatory signatures and are associated with altered patient survival

To contextualize these transcriptional changes within clinically relevant GBM phenotypes, we performed gene set enrichment analysis (GSEA) using recently defined GBM meta-programs (MPs) that reflect tumor cell identity (e.g., glial progenitor-like, astrocyte-like, neuronal-like) or cellular activity states (e.g., hypoxia or stress responses) [30]. Microgravity-grown GBM alone organoids showed significant enrichment of a ribosomal protein meta-program (MP_1_RP; P < 1*10⁻^5^), accompanied by reduced expression of hypoxia-response (MP_5_Hypoxia; P < 0.01) and mesenchymal (MP_6_MES; P < 0.05) meta-programs (**Figure 4A**). Leading-edge analysis (**Figure 4B**) revealed downregulation of key genes involved in hypoxia-driven metabolic reprogramming (*BNIP3, SLC2A3, NAMPT, ACSL3*) and stromal remodeling (*VEGFA, ANGPTL4, SERPINH1, PLOD2*), consistent with attenuated hypoxic and mesenchymal signaling. We additionally found reduced expression of the MES2 meta-program [44], in line with downregulation of ECM remodeling and angiogenesis-promoting pathways (**Supplementary Figure S3D**).

**Figure 4.**
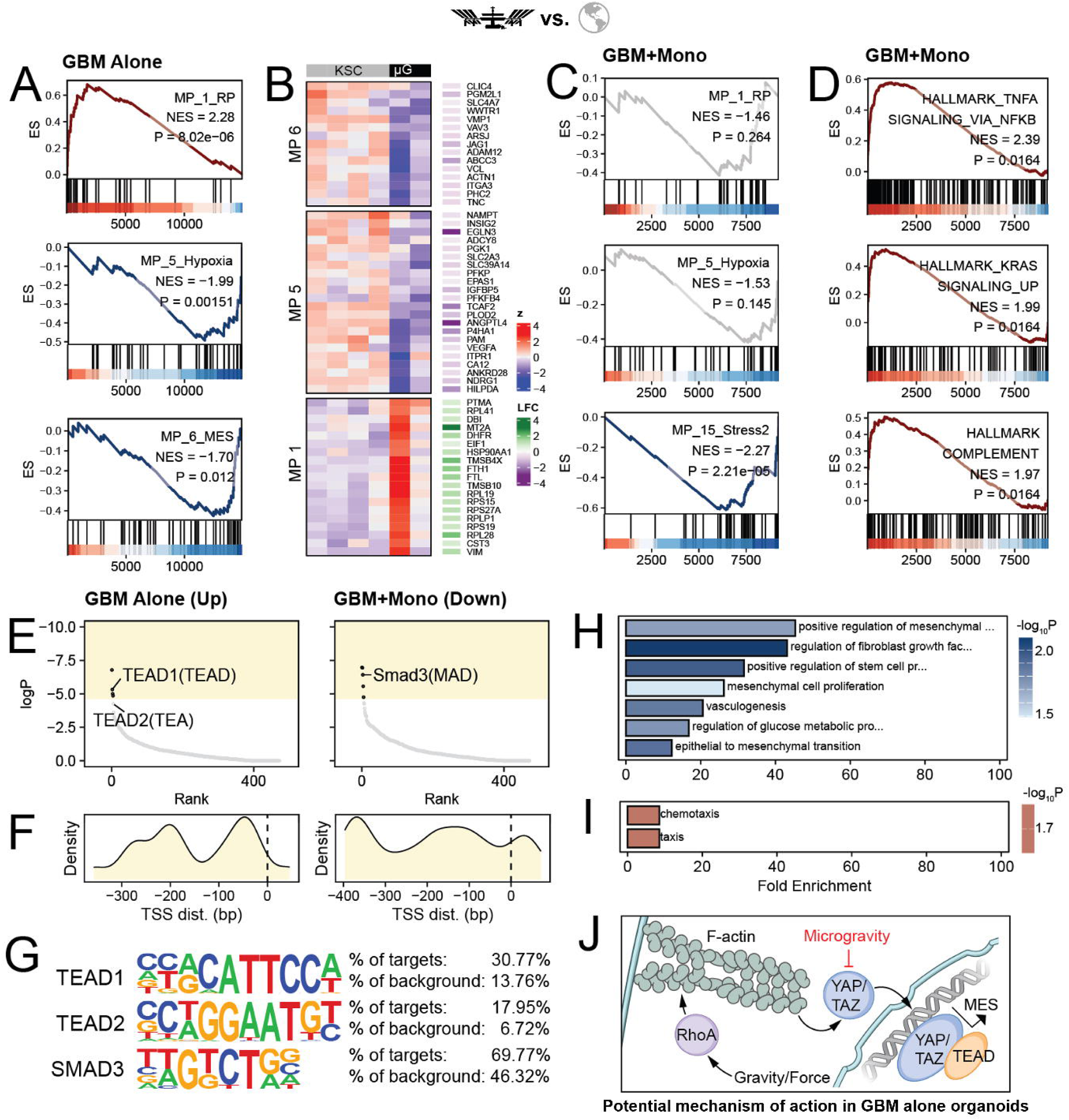
Microgravity-associated transcriptional changes align with GBM patient meta-program and TEAD/SMAD3 regulatory signatures. (**A**) Gene set enrichment analysis (GSEA) was performed using GBM meta-program (MP) gene sets [30] and Wald test DESeq2 statistic for U87 organoids grown in microgravity relative to KSC 1*g* controls. Enrichment plots are shown for MP_1_RP (ribosomal protein program), MP_5_Hypoxia (hypoxia response), and MP_15_Stress2 (stress response). In each panel, the running enrichment score (ES) is plotted across the ranked gene list (top), with tick marks indicating positions of MP genes in the ranked list (middle), and the underlying ranking metric shown below (bottom). The annotated inset reports the normalized enrichment score (NES) and Benjamini-Hochberg corrected p-value for each meta-program. (**B**) Heatmap shows leading-edge genes (i.e. core enrichment genes) from GSEA presented in (**A**). Rows correspond to leading-edge genes, and columns represent individual samples annotated by experimental condition (top bar). The main heatmap displays mean-centered and scaled variance-stabilized expression values from DESeq2. The adjacent heatmap (“LFC”) shows the corresponding DESeq2 log2 fold change (microgravity vs control) for each leading-edge gene, with color indicating effect direction and magnitude. **(C)** GSEA of GBM meta-programs (MPs) was performed as in (**A**), but using DESeq2-ranked statistics from U87 organoids co-cultured with THP-1 monocytes grown in microgravity relative to KSC 1*g* controls. **(D)** GSEA was performed on U87 organoids co-cultured with THP-1 monocytes using MSigDB Hallmark gene sets, with genes ranked by the DESeq2 Wald test statistic. (**E**) Scatter plots summarize HOMER motif enrichment results, with motifs ordered by significance (x-axis, rank) and plotted by motif enrichment log p-value (y-axis; more significant motifs appear higher due to reversed y-scale). Known motif enrichment results are shown for downregulated genes in U87 GBM (left) and upregulated genes in U87 + THP-1 monocyte co-culture (right). Each point represents one annotated HOMER motif; motifs passing significance threshold (P < 0.01, shaded region) are highlighted, and select motifs are labeled. (**F**) Density plot of HOMER-called motif offsets relative to the transcription start site (TSS) across genes used for enrichment analysis in (**I**); the dashed vertical line marks the TSS (0 bp). (**G**) Sequences logos and enrichment statistics are shown for significantly enriched motifs, based on analysis shown in (**E**). Gene Ontology (GO) over-representation analysis (ORA) was performed on downregulated and upregulated genes in U87 (**H**) and U87 + THP-1 monocyte (**I**) organoids, respectively, whose regulatory regions contained at least one TEAD1/2 (in U87; panel **H**) or Smad3 (U87+THP-1; panel **I**) motif match. Enriched GO terms were simplified to reduce redundancy. Bar plots display selected Biological Process terms (top terms by significance), with bar length indicating fold enrichment and color encoding statistical significance (-log10 p-value, Benjamini-Hochberg corrected). (**J**) Schematic demonstrating likely mechanism by which microgravity impacts GBM phenotype. Under normal gravity or mechanical force, RhoA promotes F-actin polymerization, facilitating YAP/TAZ nuclear localization and interaction with TEAD transcription factors to drive mesenchymal (MES) gene expression. In microgravity conditions, reduced mechanical loading disrupts cytoskeletal tension, leading to inhibition of YAP/TAZ activation and decreased transcription of MES-associated genes.

GBM organoids with monocytes similarly exhibited reduced expression of the stress-response meta-program (P < 1*10⁻); however, no significant changes were observed in other GBM meta-programs (**Figure 4C**). To assess inflammatory signaling in microgravity-grown GBM-monocyte organoids, we performed GSEA using the MSigDB Hallmark gene set collection. This analysis revealed enrichment of NFkB signaling, KRAS signaling, and complement pathways (**Figure 4D**).

To identify putative upstream regulators of the observed transcriptional changes, we performed motif enrichment analysis using Hypergeometric Optimization of Motif EnRichment (HOMER), focusing on regions spanning 400 bp upstream and 100 bp downstream of transcription start sites (TSSs) of up-and downregulated genes. In GBM alone organoids, genes downregulated by microgravity exposure were enriched in TEAD1 (P=0.01) and TEAD2 motifs (P=0.1), whereas genes upregulated in GBM-monocyte organoids were enriched in Smad3 motif (P=0.01) (**Figure 4E**).

We next more closely examined potential regulatory relevance of these transcription factors by mapping TEAD1/2 and Smad3 binding motifs relative to TSSs (**Figure 4F**). Consensus motif sequences were enriched within 400 bp upstream of the TSS, suggesting proximal promoter-mediated regulation (**Figure 4G**). Overrepresentation analysis of TEAD1/2 and Smad3-associated DEGs revealed repression of mesenchymal pathways (**Figure 4H**) and activation of chemotaxis (**Figure 4I**) in GBM alone and GBM-monocyte organoids, respectively.

Finally, we evaluated whether transcriptional changes induced or repressed by microgravity are associated with more aggressive tumor biology found in GBM patients by analyzing RNA-sequencing data available from The Cancer Genome Atlas (TCGA). We found a significant association between the gene signature upregulated in microgravity-grown GBM alone organoids (the top 50 DEGs based on Wald statistic) and longer progression-free survival in GBM patients, as assessed by log-rank test (**Supplementary Figure S4A**) (P < 0.01) and Cox proportional hazard model with key clinical covariates accounted for (**Supplementary Figure S4B**) (P < 0.05).

Together, these findings indicate that microgravity attenuates mesenchymal transcriptional state in microgravity-grown GBM alone organoids, possibly through the YAP/TAZ/TEAD transcriptional regulators. TEAD transcription factors are the primary DNA-binding partners through which YAP and TAZ regulate gene expression; accordingly, enrichment of TEAD1/2 binding motifs proximal to target gene TSSs, coupled with repression of mesenchymal gene expression, is consistent with reduced engagement of YAP/TAZ-driven mesenchymal programs under microgravity conditions (**Figure 4J**).

### Microgravity-induced changes in gene expression patterns vary with myeloid cell subtype

We next applied Xenium *in situ* transcriptomics on agarose-embedded organoids to resolve the spatial organization of GBM transcriptional states in microgravity-grown organoids compared to ground controls. Cell segmentation identified 6,668 cells across all samples, of which 5,239 passed quality-control filtering based on transcriptome complexity and were retained for downstream analyses.

Using the filtered dataset, we first assessed cellular composition to determine if GBM organoids co-cultured with THP-1 monocytes contained monocytes or macrophages that persisted following long-term organoid culture (regardless of gravitational conditions). To this end, we generated a single-cell reference by performing scRNA-seq profiling of U87 GBM cells and integrating these data with a publicly available scRNA-seq dataset of THP-1 monocytes ([45]; **Supplementary Figures S5A-B**). Cell identities in the Xenium dataset were inferred using robust cell type decomposition (RCTD), a Poisson mixture model that assigns each cell weights corresponding to the likelihood that its expression profile is explained by each reference population. RCTD-derived weights indicated that most cells were confidently classified as U87 GBM cells (**Supplementary Figure S5C**). Consistent with this result, we detected no expression of canonical monocyte marker genes in the Xenium dataset, with the exception of CSF1R and SIRPA, which were co-expressed by both THP-1 monocytes and GBM cells in the reference data (**Supplementary Figure S5D**). Together, these findings provide further evidence for limited persistence of THP-1 monocytes or macrophages following long-term organoid culture.

To identify dominant axes of transcriptional heterogeneity within GBM cells, we performed consensus non-negative matrix factorization (cNMF) on the Xenium dataset. NMF identifies gene expression programs (GEPs) such that the transcriptome of each cell is represented as a linear combination of multiple programs, allowing individual cells to simultaneously express varying levels of distinct transcriptional axes. Model selection based on stability and reconstruction error identified k = 4 as the most robust solution (**Supplementary Figure S6A**), supporting the existence of four major GEPs. All four GEPs were detected across samples, experimental, and gravitational conditions, indicating that they capture shared biological programs rather than sample-specific technical variation (**Supplementary Figure S6B**). Furthermore, pairwise correlations between GEPs were minimal, suggesting that the identified programs are largely orthogonal and non-redundant (**Supplementary Figure S6C**).

To place these GEPs in the context of clinically relevant GBM states, we performed GSEA using GBM meta-programs (MPs) ([30]; **Figure 5A**). GEP1 was significantly enriched for stress-response meta-programs (MP_10 and MP_15; P < 1 * 10⁻), while GEP2 showed enrichment for the mesenchymal (MES) meta-program (MP_6). GEP3 was enriched for neuronal and glial lineage programs, including the neuregulin neuron (MP_14_NRGN; P < 0.001) and oligodendrocyte progenitor-like (MP_2_OPC; P < 0.05) meta-programs. These annotations were further supported by enrichment analyses using MSigDB Hallmark and Gene Ontology gene sets (**Supplementary Figures S6D, E**). Based on these results, we annotated GEP1 as stress response, GEP2 as mesenchymal, GEP3 as inflammatory, and GEP4 as RNA processing, reflecting the major transcriptional axes present in GBM organoids (**Figure 5B, Supplementary Table S5**).

**Figure 5.**
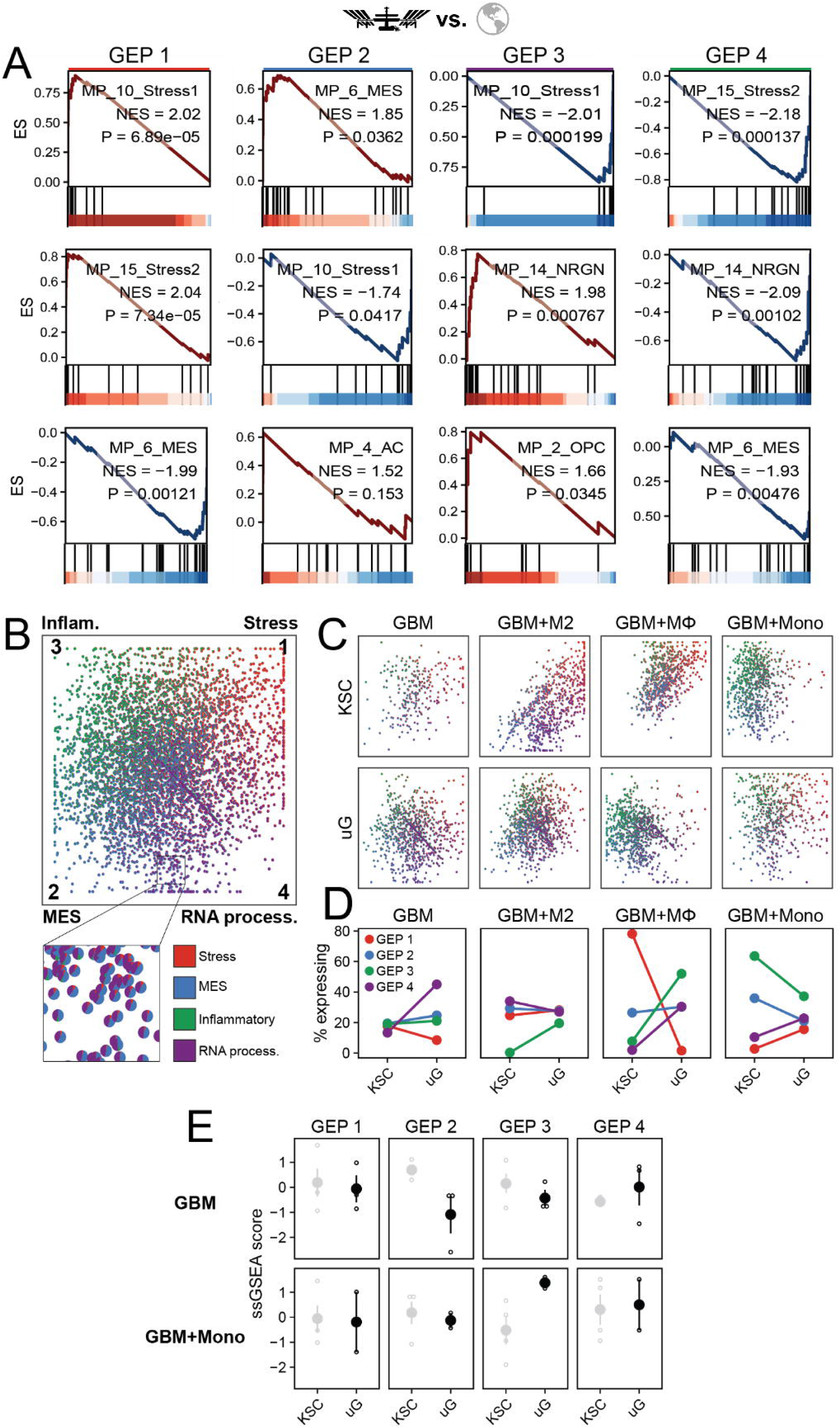
Spatial transcriptomics reveal changes in stress, mesenchymal, inflammatory, and RNA processing-associated pathways in microgravity depending on myeloid cells incorporated. (**A**) GSEA of GEPs (arranged in columns) defined by consensus non-negative matrix factorization (cNMF) was performed using GBM meta-programs (MPs, arranged in rows) [30]. Enrichment plots are shown for MP_2_OPC (Oligodendrocyte-Progenitor-like Cells), MP_4_AC (Astrocyte-like Cells), MP_6_MES (Mesenchymal-like Cells), MP_10_Stress1 (Cellular Stress Response 1), MP_14_NRGN (NRGN neuron), and MP_15_Stress2 (Cellular Stress Response 2). In each panel, the running enrichment score (ES) is plotted across the ranked gene list (top), with tick marks indicating positions of MP genes in the ranked list (middle), and the underlying ranking metric shown below (bottom). The annotated inset reports the normalized enrichment score (NES) and Benjamini-Hochberg corrected p-value for each meta-program. (**B**) Quadrant plot showing the relative usage of four gene expression programs (GEP1-GEP4) across four quadrants. The two diagonal axes represent pairwise contrasts between GEP2 vs. GEP1 and GEP3 vs. GEP4, such that cells located toward opposite ends of a diagonal predominantly express the corresponding GEP. Each point is shown as a pie chart representing a single cell, with slice areas proportional to normalized GEP usage scores. (**C**) Separate scatterpie plots are displayed for each organoid condition. KSC, Kennedy Space Center (1*g* control); uG, microgravity. (**D**) Line plots summarizing the frequency of cells assigned to each gene expression program (GEP1-GEP4) across organoid conditions. KSC, Kennedy Space Center (1*g* control); uG, microgravity. (**E**) Single-sample GSEA (ssGSEA) was used to quantify enrichment of cNMF-derived gene expression programs (GEP1-GEP4) in bulk RNA-seq samples. For each GEP, 50 top-scoring genes were used as gene sets for ssGSEA. Bulk RNA-seq expression values were variance-stabilized, ssGSEA scores were computed using GSVA, and resulting scores were mean-centered and scaled. Shown are ssGSEA scores (mean ± SE) for microgravity-grown (uG) GBM organoids with and without monocytes, together with their respective controls (KSC).

For each cell, we next determined which GEPs were significantly active by comparing observed GEP usage scores to a null distribution generated by permutation of gene expression values. Using a significance threshold of p < 0.05, we found that 19% of cells could not be confidently assigned to any GEP, 59% of cells were assigned to a single dominant GEP, and 21% of cells co-expressed two GEPs, consistent with partial state mixing often observed *in vivo* [30] (**Supplementary Figure S6F**).

We then examined how GEP usage differed between microgravity-grown organoids and ground controls (**Figures 5C, D**), focusing on changes exceeding a two-fold difference. Microgravity-grown GBM alone organoids, as well as organoids co-cultured with monocytes and macrophages, exhibited a marked increase in GEP4 (RNA processing), consistent with the upregulation of post-transcriptional

RNA regulatory pathways observed in our bulk RNA-seq analyses. GBM-macrophage organoids showed a pronounced reduction in GEP1 (stress response) accompanied by an increase in GEP3 (Inflammatory), while organoids with M2-like macrophages displayed an increase in GEP3 only.

To determine if transcriptional changes observed in bulk RNA-seq could be explained by shifts in GEP activity under prolonged microgravity exposure, we used GEP gene signatures to score bulk mRNA-sequencing datasets from GBM organoids cultured with and without monocytes. This analysis revealed trends consistent with the spatial data, including reduced expression of the mesenchymal GEP (GEP2) in GBM alone organoids and increased expression of GEP3 (Inflammatory) in organoids with prior monocyte exposure (**Figure 5E**), supporting the interpretation that bulk transcriptional changes reflect altered contributions of the underlying GEPs.

### Spatial transcriptomic analysis reveals microgravity-dependent spatial patterning of GBM cell states

We next investigated the spatial organization of GEPs within the organoids by quantifying spatial autocorrelation of per-cell GEP scores using Moran’s I. Among the four programs, GEP2 showed a trend toward increased spatial autocorrelation in microgravity-grown organoids, with more modest effects observed for other GEPs (**Figure 6A**). To assess whether microgravity globally enhances spatial clustering of transcriptional states independent of organoid composition or GEP identity, we fit a linear model across all conditions, which revealed a statistically significant effect of microgravity on spatial autocorrelation (β = 0.0895, p = 0.0183).

**Figure 6.**
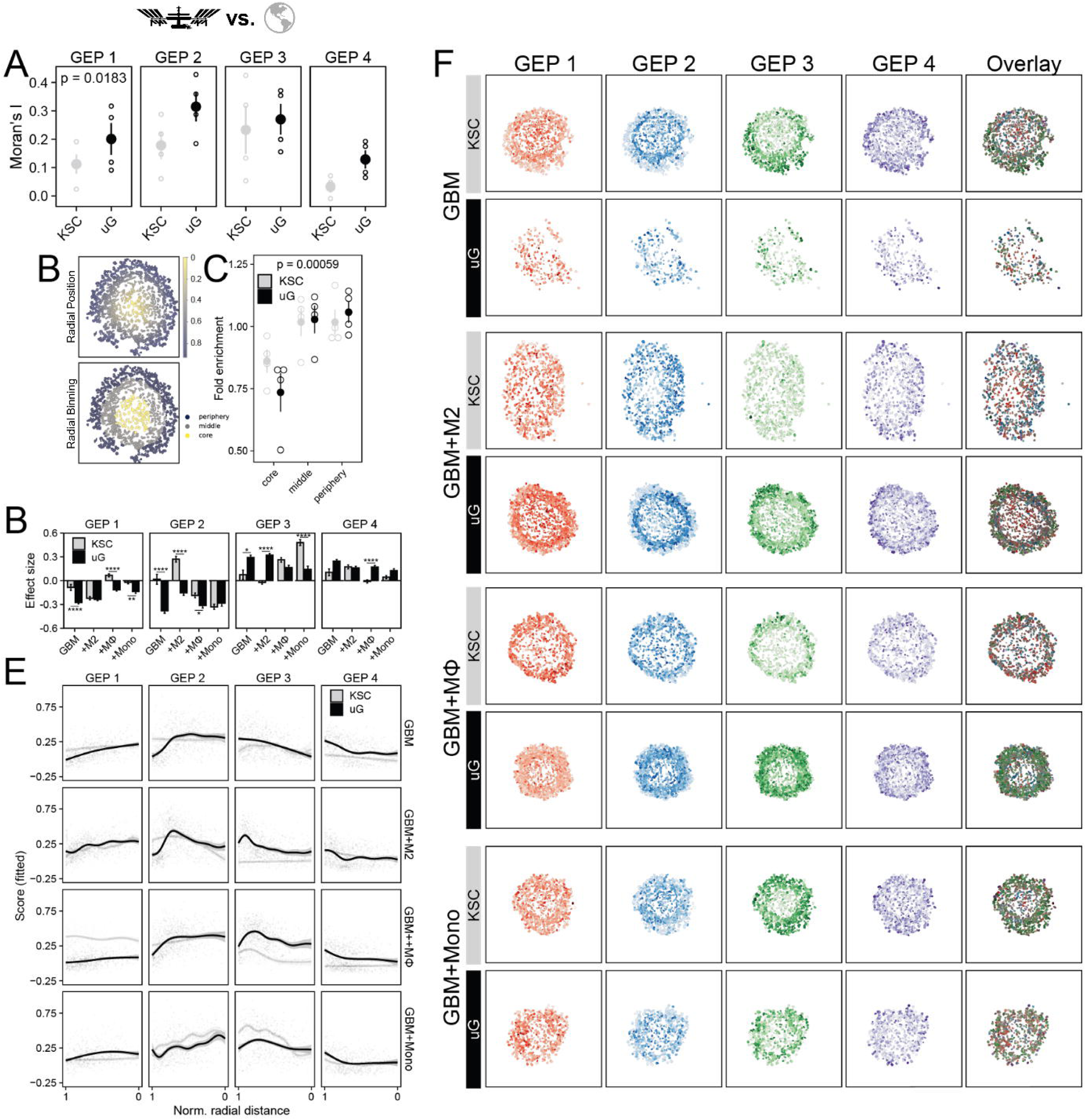
Microgravity induces distinct changes spatial patterning of gene expression patterns depending on organoid composition. (**A**) Spatial autocorrelation of GEP scores was quantified using Moran’s I for each sample. The figure shows Moran’s I mean ± SE for microgravity-grown organoids (uG) and controls (KSC). (**B**) Cells were assigned a normalized radial position defined as the cell centroid distance relative to the reconstructed organoid boundary divided by the maximum interior distance (1 = boundary, 0 = deepest interior). In the upper subpanel, cells are colored by normalized radial distance. Cells were then discretized into three equal-depth radial bins using fixed cutoffs: periphery (1-2/3), middle (2/3-1/3), and core (1/3-0). Cells with undefined radial position (outside the organoid boundary) were not considered. In the lower subpanel, cells are colored by radial bin (periphery/middle/core). (**C**) For each radial bin, observed cell counts were compared to the expected counts under uniform cell density, computed from the measured area of each bin (expected = total_cells * layer_area / total_area). Fold enrichment was quantified as the observed-to-expected ratio. Values >1 indicate increased cell density and values <1 indicate depletion. Shown are fold enrichment scores for (mean ± SE) for microgravity-grown (uG) and control (KSC) organoids. p value was calculated by testing whether the region-adjusted mean fold enrichment for each radial bin differed from 1 using a linear model and estimated marginal means (two-sided Wald test). (**D**) To quantify radial organization of GEPs, GEP scores were modeled as a function of normalized radial distance relative to the organoid boundary using linear regression. For each organoid condition and each GEP, an interaction model was fit. From each fitted model, perturbation-specific radial gradients were estimated as the marginal slopes of GEP score with respect to radial distance using estimated marginal trends (emtrends). Bars represent the estimated radial slope (effect size) ± SE. Statistical significance reflects pairwise contrasts of radial slopes between uG and KSC; p values were adjusted for multiple testing and significance is indicated by asterisks (* p < 0.05, ** p < 0.01, *** p < 0.001, **** p < 0.0001). (**E**) To characterize nonlinear radial gradients in GEP activity, per-cell GEP scores were modeled as smooth functions of normalized radial distance using generalized additive models (GAMs). For each GEP (GEP1-GEP4), models were specified with condition-specific smooth terms. Model predictions were evaluated on a dense grid of radial positions spanning the 1st to 99th percentiles of the observed radial-distance distribution. Fitted smooths are shown with 95% confidence intervals. Raw single-cell GEP scores are overlaid as semi-transparent points to indicate raw data distribution. For visualization, the radial distance axis is linearly rescaled such that the plotted domain spans 1-0. (**F**) Cell boundary polygons colored by GEP scores. Field-of-view is standardized across all samples using a single global span.

Motivated by the increased spatial autocorrelation, we next examined radial patterning of GEPs within organoids. For each cell, we computed a normalized radial distance from the organoid center, a metric that reflects gradients in oxygen and nutrient availability within the static culture system that is diffusion-dominated (**Figures 6B**). Consistent with this interpretation, we observed a significant reduction in cell density within the organoid core relative to peripheral regions (p < 0.001), with a trend toward further reduced core cell density in microgravity-grown organoids, consistent with a more hypoxic core environment (**Figure 6C**).

To quantify monotonic radial trends in GEP activity, we modeled per-cell GEP scores as linear functions of normalized radial distance. Across conditions, GEP1 and GEP2 generally increased toward the organoid core, whereas GEP3 and GEP4 decreased (**Figure 6D**). With the exception of GEP3 in organoids with monocytes and GEP2 in organoids with M2-like macrophages, radial gradients were consistently steeper in microgravity-grown organoids, indicating that microgravity enhances spatial organization of GBM transcriptional states.

Finally, to capture potential nonlinear spatial effects, we modeled GEP activity as smooth functions of radial distance using generalized additive models (GAMs) (**Figures 6E-F**). In GBM alone organoids, the most pronounced differences were observed in the organoid periphery, where microgravity-grown samples exhibited higher GEP3 and GEP4 activity alongside reduced GEP1 and GEP2 activity, with these differences diminishing toward the organoid core. Similar region-specific effects were observed for GEP2 and GEP3 in organoids with M2-like polarized macrophages and monocytes, respectively. All other microgravity-associated changes in GEP activity were spatially uniform, indicating global rather than region-specific effects.

### Microgravity alters the cytokine secretome of GBM + monocyte organoids

Conditioned medium from the microgravity and ground control organoids was collected immediately upon sample retrieval for proteomics analysis. These samples were analyzed using Olink’s high-sensitivity proximity extension assay with a panel of 45 human cytokines to obtain absolute protein concentrations. GBM-monocyte organoids exhibited significant changes in the secretion of six cytokines (**Figure 7A**). Microgravity-grown organoids secreted more CXCL12, which is elevated in GBM tissues and is implicated in GBM progression, invasion, treatment resistance, and recruitment of pathological myeloid cells [46], [47], [48]. Elevated CXCL12 in tumors of glioma patients is correlated with more rapid tumor recurrence [49]. LOX-1 was also more abundant in microgravity, and contributes to tumor progression, angiogenesis, invasion, and immune modulation and is predictive of shorter overall survival in GBM patients [50], [51], [52]. The cytokines IL-13 and IL-17A are both implicated in GBM migration, invasion, and proliferation, and were increased in microgravity compared to ground controls [53], [54]. IL-27, also increased in microgravity, has pleiotropic roles in cancer, potentially contributing to tumor suppression or to immune suppression, depending on the context [55]. CSF3 alone was decreased in microgravity, and is associated with immune cell recruitment [56]. Two additional cytokines, IL33 and CSF2, reflected modest changes in overall abundance, but may have differences in their spatial expression within the organoid not captured by bulk secretome analysis (**Figure 7B, C**). IL33 gene expression was relatively low at the organoid core and middle but was significantly elevated towards the periphery (**Figure 7B**). CSF2 expression, in contrast, was highest in the organoid core and decreased in the middle and periphery (**Figure 7C**).

**Figure 7.**
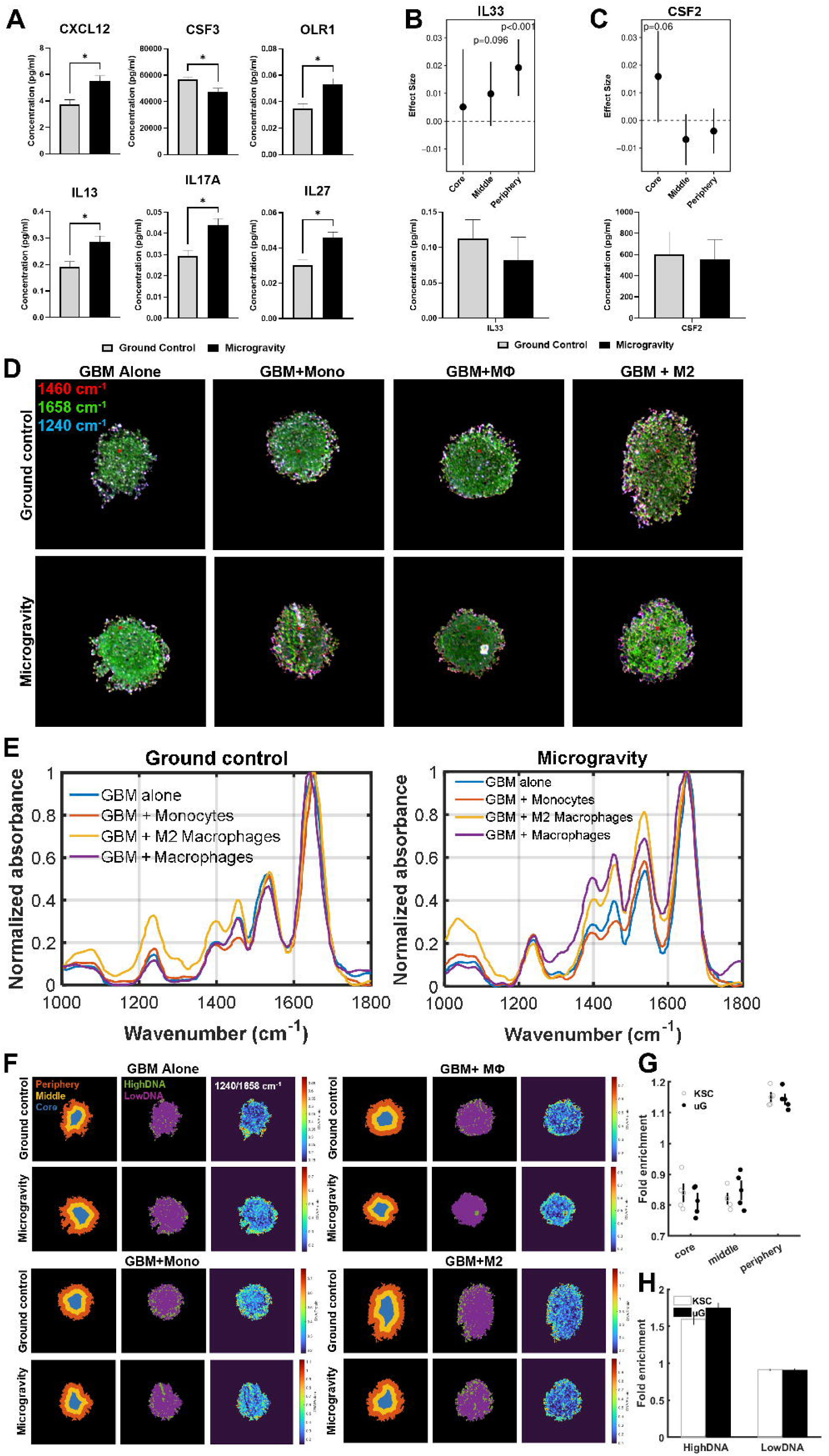
Microgravity induces alterations in protein secretome and spatial spectral profiles of GBM organoids. (**A**) Quantification of concentration of cytokines in GBM+monocyte conditioned media via the Olink^®^ Proximity Extension™ Assay Target48 cytokine panel. Statistical comparisons were made using unpaired t tests (* p < 0.05). Error bars indicate SEM. Plots of spatial gene expression of IL33 (**B**) and CSF2 (**C**) in three radial bins with all organoid conditions combined (upper) and corresponding media concentration (lower) of combined samples. p value was calculated using estimated marginal means for experimental condition (microgravity vs KSC) conditioned on radial layer using linear interaction model. (**D**) Images of each organoid reconstructed as a pseudo-RGB image from spatial FTIR data absorbance measurements. Images show absorbance at 1460 cm^-1^ (red), 1658 cm^-1^ (green), and 1240 cm-1 (blue). (**E**) Representative DF-IR spectra from ground control and microgravity across all organoid onditions. The yellow x on the pseudo-RGB image marks the pixel used to extract the representative point spectrum. (**F**) Illustration of spatial binning of each organoid into core, middle, and periphery regions (left column), images colored for regions of high or low DNA content (middle column), and heatmap images of 1240/1658 cm^-1^ ratios for each organoid (right column) for ground control and microgravity organoids. Arrows indicate regions of spectral difference between ground control and microgravity. (**G**) The calculated enrichment scores for high DNA content in the core, middle, and periphery of ground control and microgravity organoids. (**H**) Enrichment of HighDNA and LowDNA regions in microgravity versus ground controls.

### Microgravity-induced and spatial biochemical trends are detectable by chemical imaging

We next performed quantum cascade laser-based discrete frequency infrared (DF-IR) imaging on FFPE organoid sections, obtaining absorbance values of bands associated with Amide I, lipid/protein content, ECM, nucleic acids, and phospholipids. Images of these bands, normalized to respective control bands, are shown in **Supplementary Figure S7A.** Overlay images of the 1460, 1658, and 1240 cm⁻¹ images (corresponding to lipid/protein content, Amide I, and nucleic acids) are shown in **Figure 7D**. Microgravity-grown organoids tended to have higher absorbance compared to ground control in these fingerprint regions, particularly in Amide I and nucleic acid (**Figure 7E**). To evaluate radial trends, each organoid image was segmented into core, middle, and periphery regions (**Figure 7F**). The 1240/1658 cm⁻¹ ratio (nucleic acids) showed a clear increase at the periphery compared to the middle and core (**Figure 7G**), consistent with the cell density data (**Figure 6C**), implying the formation of a necrotic core. However, there was no significant difference in the amount of DNA-rich area between ground control and microgravity organoids (**Figure 7H**). For the Beta/Alpha proxy, ECM/Protein, Lipid/Protein, and Phosphate/Protein ratios, there was also a trend towards increased expression at the periphery compared to the core regions while the Amide I/Amide II ratio was slightly reduced at the periphery, with similar trends for both microgravity and ground control organoids (**Supplementary Figure 7B-G**).

Organoid composition tended to alter the direction of chemical response to microgravity. GBM+M2 macrophage and GBM+macrophage organoids were the most distinct, with unpolarized macrophages having decreased DNA, Lipid, Phosphate, and ECM signals in microgravity, and M2-like macrophages inducing an increase in those species, across the core, middle, and periphery regions (**Supplementary Figure S8**). These observations suggest that microgravity does not elicit a uniform global shift in spectral intensity. Rather, it may induce immune-context–dependent shifts in vibrational band ratios.

## DISCUSSION

Here, we report the first published investigation of long-term GBM organoid growth and development in space. As others have shown, spaceflight can facilitate models of accelerated disease, spontaneous spheroid formation, and immune dysregulation, and is thus a valuable tool to gain insight into diseases on Earth [4], [5], [6], [57], [9], [33], [58]. Work published to date on the effects of microgravity on GBM is limited to either ground-based simulated microgravity models or short-duration suborbital flight, both of which involve significant mechanical perturbations (e.g., fluid flow and vibrations in simulated microgravity devices, hypergravity during takeoff and landing, etc.) that may dwarf the relative contribution of actual time spent in reduced-gravity conditions [3], [14]. Utilizing the high-throughput, low-maintenance, and robust CubeLab system for automated long-term static culture, we sent an array of GBM-myeloid organoids to the ISS as part of the SpX-30 mission. After 40 days, the organoids grown in microgravity had superior morphological and transcriptional organization, with the inclusion of myeloid cells contributing to the development of transcriptional markers of a more advanced, aggressive GBM model.

We first established that our GBM-immune organoids can be maintained in sealed cryovials for long-term culture, reaching a steady state featuring a necrotic core surrounded by live cells, and an acidified and likely hypoxic media environment, mimicking what is observed in GBM tumors [36]. Monocytes or macrophages were added to these organoids, showing uniform incorporation in the initial micro-organoids. Interestingly, organoids incorporating M1-like polarized macrophages were much less likely to remain intact and grow. This suggests the successful induction of an antagonistic interaction between the pro-inflammatory macrophages and the tumor cells, consistent with similar reports that polarized microglia induce death in glioma organoids [59]. Morphological analysis of the organoids after 40 days in space confirmed our hypothesis that microgravity promotes the formation of more uniform and compact organoids, giving them an advantage over Earth-based models that are subject to settling and disaggregation.

GBM alone organoids featured enrichment of genes associated with post-transcriptional regulation, and a reduction in genes linked to mesenchymal-like behaviors such as ECM organization and angiogenesis, glycolysis, and stress response. The gene signature that was upregulated in microgravity by GBM alone organoids was associated with better patient survival, consistent with studies of GBM cells in simulated microgravity, which report reduced proliferation and invasion and increased apoptosis and chemosensitivity [60], [61], [62], [63], [64], [65]. In contrast, GBM+monocyte organoids upregulated gene sets and individual genes associated with detrimental tumor traits such as chronic inflammation and tissue and vascular remodeling, leading to GBM progression, treatment resistance, and immune dysfunction [39], [40], [41], [42], [43]. For example, they had increased activity of TNFα signaling via NF-kB and KRAS signaling, which are highly active in GBM tumors and influence tumor progression, invasion, inflammation, and response to treatment [66], [67]. KRAS signaling plays a critical role in gliomagenesis and tumor maintenance, and has also been observed to upregulated in astronaut skin biopsies after spaceflight [68], [69]. There was little overlap between the space-associated DEGs in GBM alone and GBM+monocyte organoids identified from bulk RNA sequencing. This is consistent with prior observations that the effects of spaceflight is cell-type specific, and that a single-cell analysis approach is necessary to identify changes which could be missed in bulk RNA sequencing [13].

Therefore, we next performed spatial transcriptomics analysis on a panel of 5,000 genes using Xenium’s 10X platform. From this single-cell level dataset, we identified four distinct GEPs corresponding to stress response, mesenchymal state, inflammation, and RNA processing. Microgravity induced a significant increase in GEP spatial autocorrelation across the combined organoid conditions and GEPs. Microgravity-grown organoids also had stronger radial-periphery patterning, with generally steeper enrichment gradients between the core and periphery. Spatial analysis of cell states of glioma patient samples show that a mesenchymal-hypoxia state is generally highly spatially organized, and that regions of the tumor expressing this program are surrounded by a layer of inflammatory and immune-associated cell states [28]. Mimicking this, our GBM alone and GBM+M2-like macrophage organoids grown in microgravity upregulated the mesenchymal program in the core, surrounded by an inflammatory program-enriched periphery. Thus, microgravity enhances the spatial organization of these key cell states to more closely mimic what is observed in patients.

Proteomics analysis provided further support the role of microgravity in the formation of a well-developed, immunosuppressive organoid microenvironment. This was evidenced by the upregulation of secreted cytokines implicated in GBM progression, invasion, treatment resistance, and immune evasion. Finally, DF-IR imaging confirmed the spatially distinct biochemical composition [70] of the core, middle and periphery of the organoids, including nucleic acid-associated phosphate vibrational modes, aliphatic vibrational modes indicative of lipid/protein composition, and protein concentration and conformation-sensitive amide I/II features. Laboratory-based laser scanning microscopes [71], [72] have only recently attained the theoretical performance [73] previously demonstrated exclusively on synchrotron-based Fourier transform infrared imaging systems [74]. To our knowledge, this study represents the first instance of using IR chemical imaging to interrogate spatial biochemical remodeling in organoids grown under microgravity conditions. Further study is warranted to validate the observed trend towards immune-dependent differences in chemical response to microgravity.

While we see a clear difference in the effects of microgravity when monocytes were present compared to GBM alone conditions, bulk analysis and single-cell spatial analysis leads us to conclude that monocytes were unlikely to have persisted to the experiment endpoint regardless of gravitational exposure. This is unsurprising, given the static culture conditions that are likely more detrimental to immune cells than adaptive cancer cells. However, the early incorporation of monocytes and macrophages did impact response to microgravity, suggesting that even their temporary “preconditioning” role influences the disease model. Given their essential role in GBM disease progression and their potential as a therapeutic target [75], [16], [76], ongoing efforts are directed at optimizing an accurate and persistent GBM immune microenvironment in these models, which will likely produce more aggressive, immunosuppressive, and treatment-resistant organoids and will contribute to the growing field of “astroimmunology” [21].

As with any biological study conducted on the ISS, there are several technical limitations to this work. While the ground control samples were created from the same batch of organoids and transported together to KSC, this did not control for factors such as mechanical perturbations during launch, docking to the ISS, and return to Earth. The increase in success rate of agarose-embedded organoids compared to those that were free-floating in media suggests that adding a hydrogel matrix around the organoid may serve to cushion it from vibrations or impact forces. Alternatively, spatial confinement and the resulting compressive solid stress may contribute to macrophage polarization towards a tumor-supporting phenotype, as suggested by our earlier mechanobiology work [35], [77]. This highlights the need for further studies on spaceflight-grown GBM-immune organoids in brain-mimicking scaffolds to better recapitulate *in vivo* biology.

While on the ISS, the experimental payload was also exposed to increased levels of radiation, which poses an important health risk to astronauts [78]. However, the amount of radiation that these organoids experienced, approximately 8.1 mGy total, was far below the levels reported to cause measurable effects on human cells. Studies measuring the effects of simulated GCR on blood cells report minimal changes in chromosomal aberrations at their lowest tested dose, 0.1 Gy, which is more than 10 times higher than the estimated cumulative dose of our samples over the entire 40-day period in space [79], [80]. Monocytes are particularly radiation-resistant, with a minority of cells responding to doses more than 200 times higher than what our samples experienced [80]. Similarly, GBM cells have a modest reduction in viability at a dose more than 20 times higher than our samples experienced [81].

In conclusion, the present study characterizes a novel, advanced GBM organoid model leveraging the unique microgravity environment of space to better recreate the tumor immune microenvironment of advanced disease. To our knowledge, this is the first demonstration of GBM organoids in a long-duration space mission and the first spatial transcriptomics analysis of organoids generated in space. Future payloads will include patient-derived GBM organoids, additional immune subtypes, biomechanical design considerations, high-throughput drug screening, and expanded orthogonal and integrated dataset production and analysis from the organoids. The in-space organoid growth and analytical pipeline developed here will serve as the foundation for future tumor modeling and drug testing on the ISS, commercial space stations, and free flying platforms, not only for GBM but for other cancers as well. Space biomedical research and development will continue to expand to support orbital oncology efforts and capabilities to develop paradigm-shifting knowledge and therapies for humans on Earth.

## Supporting information

Table S1 Organoid Conditions

Table S2 Radiation Data

Table S3 Success Rate

Table S4 Bulk DEGs

Table S5 GEP Signature Genes

## ACKNOWLEDGEMENTS

Funding for this work was provided by the U.S. National Science Foundation and Center for the Advancement of Science in Space (CMMI: 2425684), the Air Force Office of Scientific Research Broad Agency Awards (FA9550-24-1-0456), and the Air Force Office of Scientific Research Young Investigator Program (FA9550-25-1-0113) to M.D.; Chan Zuckerberg Biohub to R.B.; and the Walther Cancer Foundation Interdisciplinary Interface Training Project Grant, the Edison Innovation Fellowship, and the Azenta Oncology Insights Grant Program (to A.B.D.). Funding for the development of Space Tango’s High Sample Capacity CubeLab was provided by the U.S. National Science Foundation SBIR Phase I Grant Program (216101) to Space Tango. We thank Ms. Melissa Stephens, Mr. Brent Harker, and Ms. Jacqueline Lopez and the Notre Dame Genomics and Bioinformatics Core Facility for performing and assisting with RNA sequencing and Xenium spatial transcriptomics. We thank Ms. Sarah Chapman at the Notre Dame Integrated Imaging Facility Histology Core for helping to optimize FFPE protocols and preparation of organoids for histological and spatial transcriptomics analysis. We thank 10X Genomics for their technical assistance with our transcriptomics and Azenta Life Sciences for their generous support of our proteomics analysis project. We thank Ms. R’nld Rumbach for technical support. We thank Mr. Fionn Lay and Ms. Anya Zhao for their assistance in experimental optimization. We thank Prof. Katharine White for generously sharing her laboratory space for experimental work. We thank Dr. Livia Luz and Prof. Alysson Muotri for experimental advice. This experiment was made possible by the International Space Station National Laboratory, the National Aeronautics and Space Administration, and Space Exploration Technologies Corp.; we thank all staff and crew involved. Figure schematics were generated using BioRender.

## AUTHOR CONTRIBUTIONS

A.B.D.: Investigation, methodology, formal analysis, visualization, writing – original draft.

M.Z.: Formal analysis, data curation, visualization, writing – original draft, writing – review & editing.

S.G. and T.C.: Methodology, project administration, and writing - review & editing. J.R., P.K.: Methodology.

K.B.: Investigation, visualization, formal analysis, writing – original draft. J.N.: Investigation and visualization.

R.B. Project administration and writing – review and editing.

M.D.: Conceptualization, methodology, project administration, funding acquisition, writing - review & editing

---

**Supplementary Figure S1.**
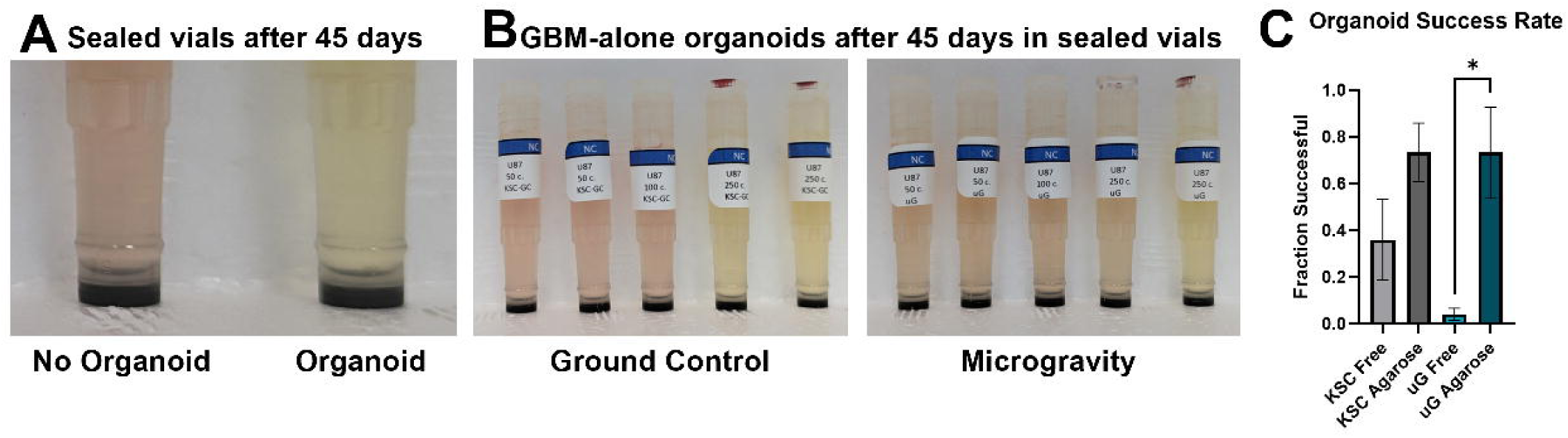
Organoid growth causes visible changes in media pH. (**A**) Image of a cryovial with no successful organoid formation with the original pink media color (left) and a cryovial containing a 45-day old organoid with acidified, yellow media (right). (**B**) Images of ground control and microgravity cryovials initially loaded with organoids made with 50 cells, 100 cells, and 250 cells. Vials on the left of each image (50-cell organoids) were more pink, indicating failure to support a successful organoid, and vials on the right (250-cell organoids) were more likely to be yellow, indicating organoid growth and media consumption. (**C**) Fraction of cryovials in each condition found to contain a successful organoid at the 45-day experiment endpoint. Error bars represent SEM. Significance was calculated with Welch’s t test, * p < 0.05.

**Supplementary Figure S2.**
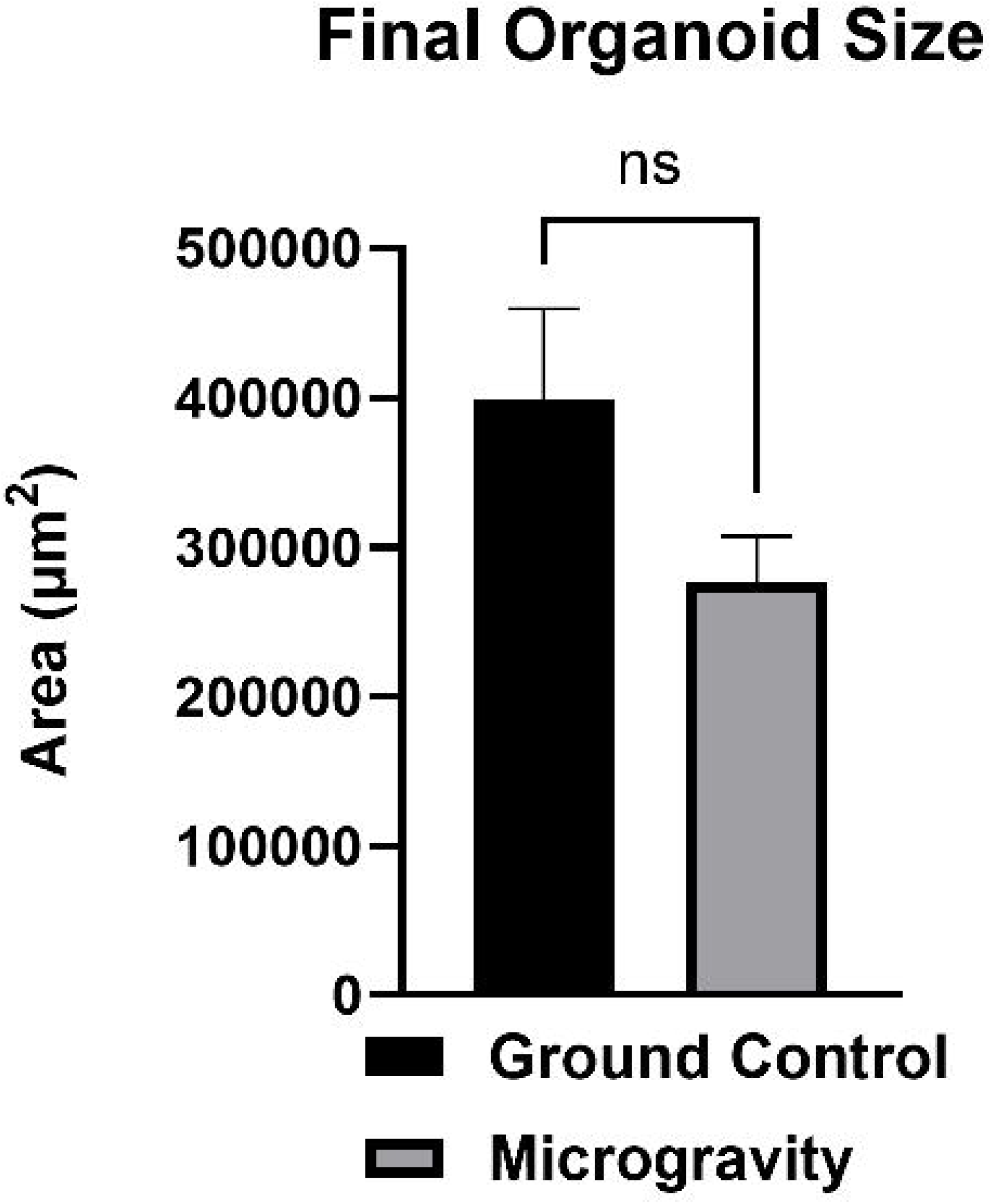
Organoids grown in microgravity have no significant difference in size. Area of ground control and microgravity-grown organoids based on brightfield images of agarose-embedded organoids. p = 0.17 via unpaired t test.

**Supplementary Figure S3.**
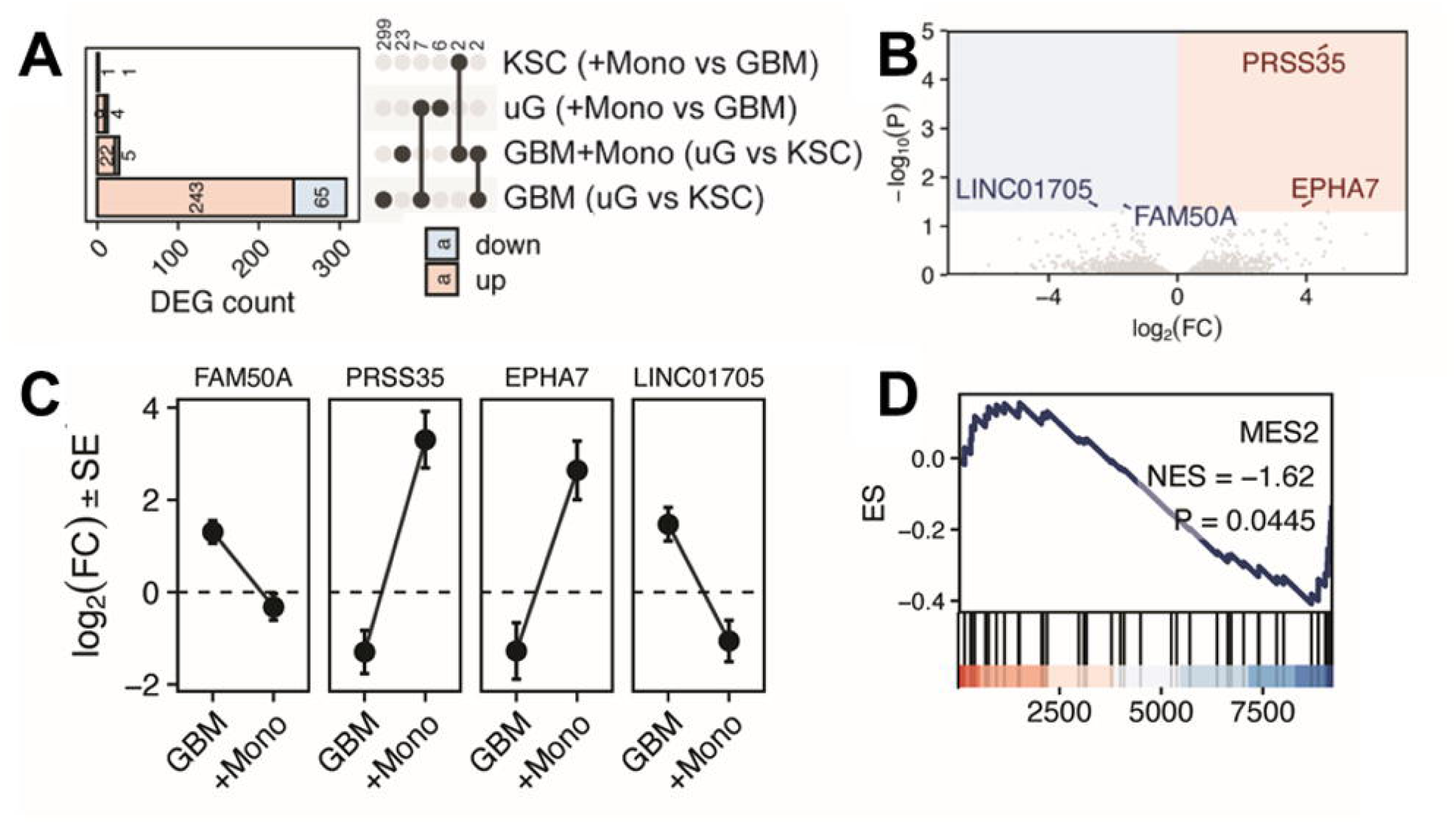
Immune context-dependent transcriptional responses to microgravity in GBM organoids. (**A**) Stacked bar plots summarize the number of significantly DEGs identified by DESeq2 (Benjamini-Hochberg corrected p < 0.05) for four contrasts derived from a model including gravity condition (uG vs KSC), co-culture condition (U87 GBM with and without THP-1 monocytes), and their interaction. Bars show DEGs for (i) U87 (uG vs KSC), (ii) U87THP (uG vs KSC), (iii) uG (U87THP vs U87), and (iv) KSC (U87THP vs U87). Counts are partitioned into upregulated (positive log2FC; light red) and downregulated (negative log2FC; light blue) genes, with the number of DEGs shown. An adjacent UpSet plot displays intersections among DESeq2-identified DEGs. Vertical bars indicate the number of genes in each intersection, and filled dots denote the contrasts contributing to each intersection. (**B**) Volcano plot shows DESeq2 results for the gravity * co-culture interaction term from a DESeq2 model including gravity condition (uG vs KSC), co-culture composition (U87 GBM with and without THP-1 monocytes), and their interaction. Each point represents a gene, plotted by the interaction log2 fold-change (x-axis; difference in the microgravity effect between U87THP and U87) and -log10 adjusted p-value (y-axis; Benjamini-Hochberg corrected). Genes with p < 0.05 are highlighted as significant (red/blue by direction), while non-significant genes are shown in gray; shaded regions indicate the significance threshold. All significant interaction genes are labeled. (**C**) Dot plots show DESeq2 log2 fold-change (uG vs KSC) estimates for selected genes (FAM50A, PRSS35, EPHA7, LINC01705) in U87 GBM alone and GBM co-cultured with THP-1 monocytes. Points indicate the estimated log2FC for each contrast, and error bars represent ±1 standard error (lfcSE). Lines connecting the two contrasts are shown to emphasize differences in direction and magnitude of the microgravity response depending on immune context. A dashed horizontal line marks no change (log2FC = 0). (**D**) GSEA of U87 GBM differentially expressed genes (uG vs KSC) was performed using GBM meta-programs (MPs) [44]. Enrichment plot is shown for MES_2. Running enrichment score (ES) is plotted across the ranked gene list (top), with tick marks indicating positions of MES_2 genes in the ranked list (middle), and the underlying ranking metric shown below (bottom). The annotated inset reports the normalized enrichment score (NES) and Benjamini-Hochberg corrected p-value for MES_2 MP.

**Supplementary Figure S4.**
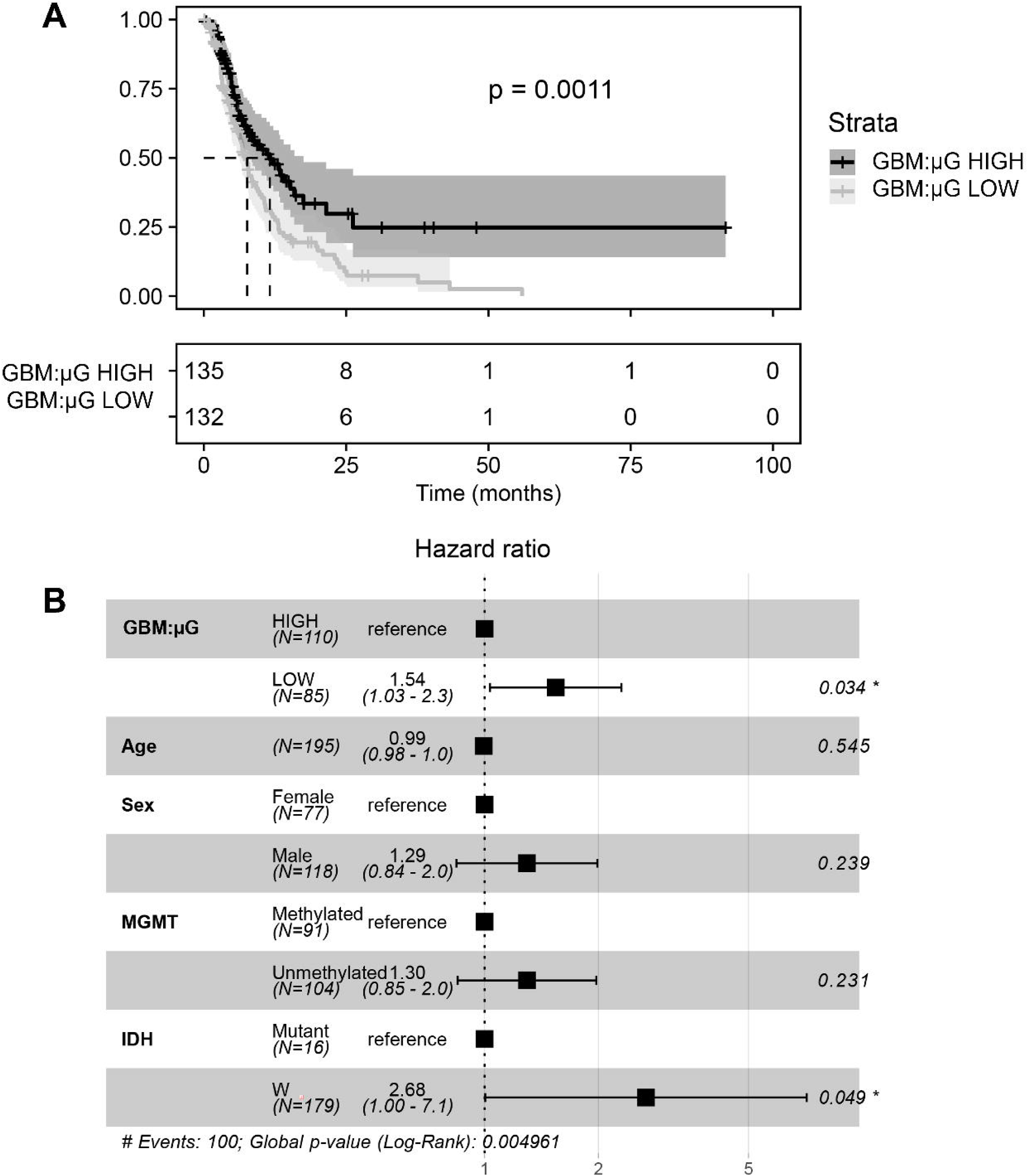
Gene signature upregulated in GBM alone organoids in microgravity is associated with improved patient survival. (**A**) TCGA RNA-seq data was used to stratify GBM patients based on single-sample gene set enrichment scores (ssGSEA) from variance-stabilized expression data using top 50 genes upregulated in the GBM alone microgravity condition based on Wald statistic. Scores were mean-centered and scaled, averaged per patient and stratified into “high” vs “low” groups using the median for survival comparison. (**B**) Associations with overall survival and progression-free survival based on a multivariable Cox proportional hazards model including the ssGSEA signature and covariates (age, sex, MGMT promoter status, and IDH status).

**Supplementary Figure S5.**
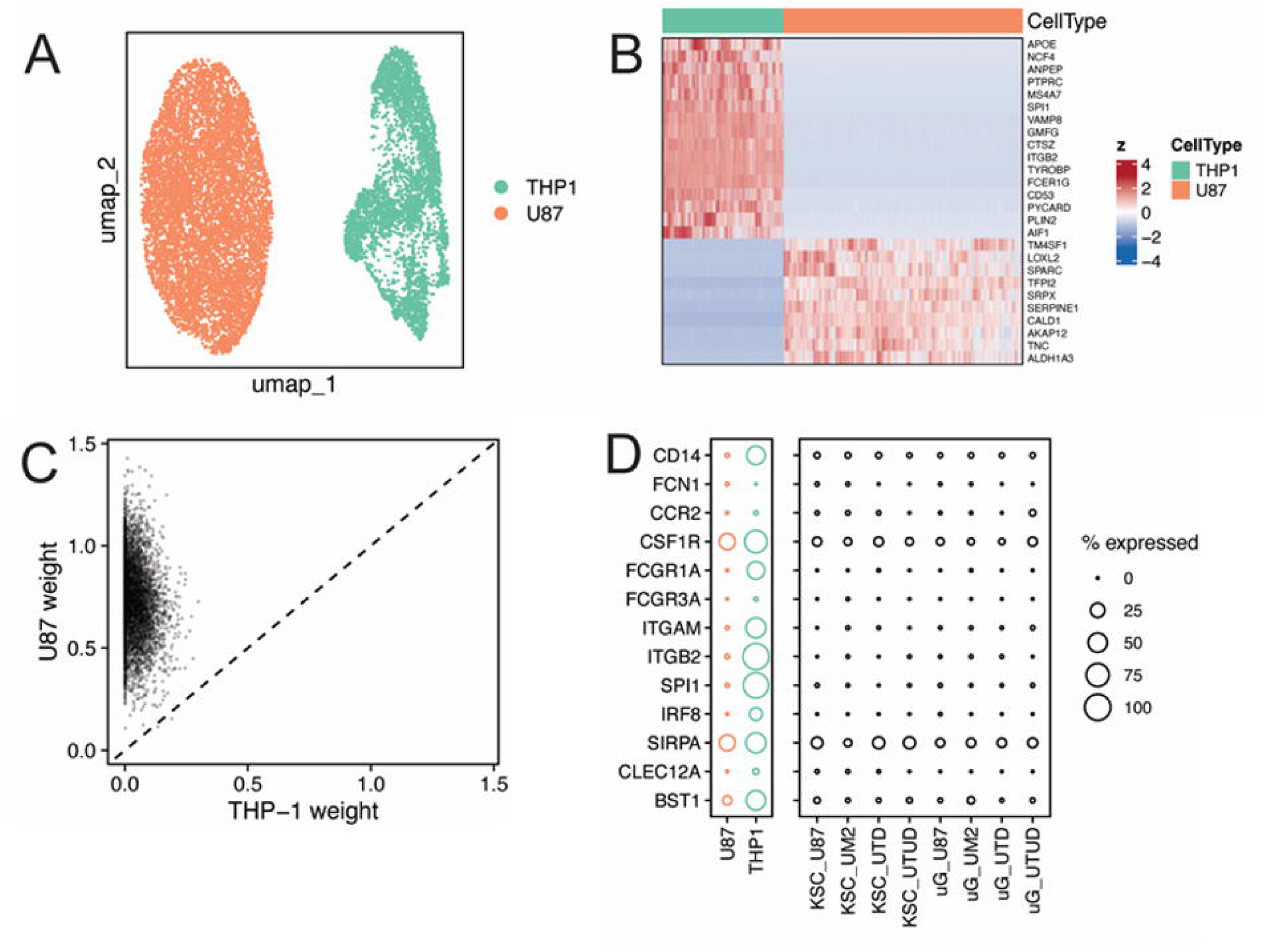
Reference-based mapping of tumor and monocyte identities to Xenium single-cell profiles demonstrates unlikely persistence of monocytes after prolonged static culture. (**A**) Uniform Manifold Approximation and Projection (UMAP) embedding of the scRNA-seq reference dataset combining U87 GBM cells and THP-1 monocytes. Cells are colored by cell type. (**B**) Heatmap of marker gene expression in the scRNA-seq reference dataset. Marker sets include myeloid/innate immune genes (e.g., TYROBP, PTPRC, FCER1G, AIF1, APOE) and mesenchymal-like GBM genes (e.g., ALDH1A3, SERPINE1, TNC, SPARC, LOXL2). Genes were selected as markers identified by Wilcoxon rank-sum test and combined into a curated, nonredundant list for visualization. (**C**) Scatterplot of cell-type weights inferred by robust cell type decomposition (RCTD). Each point represents one cell, plotted by its inferred THP-1 weight (x-axis) and U87 weight (y-axis) from the RCTD weights matrix. The dashed diagonal indicates equal weights (y = x). (**D**) Dot plots summarizing the fraction of cells expressing canonical monocyte markers (CD14, FCN1, CCR2, CSF1R, FCGR1A, FCGR3A, ITGAM, ITGB2, SPI1, IRF8, SIRPA, CLEC12A, BST1). Left: scRNA-seq reference (U87 + THP-1). Right: Xenium single-cell dataset.

**Supplementary Figure S6.**
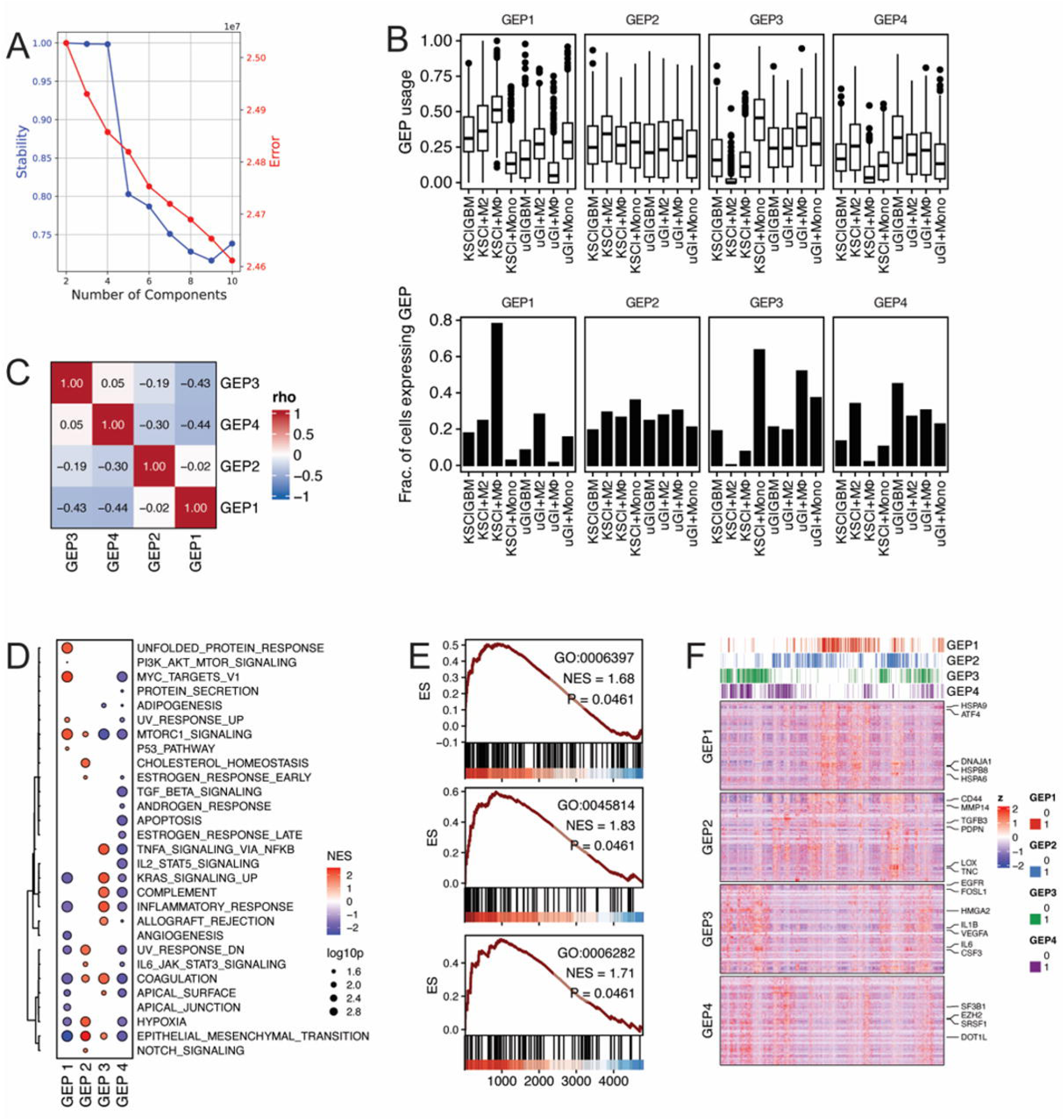
Identification and characterization of cNMF-derived gene expression programs in Xenium data. (**A**) To select the number of components for consensus non-negative matrix factorization (cNMF), cNMF was run on the merged Xenium dataset using 50 NMF iterations per candidate model. Models were fit across a sweep of component numbers (K = 2-10). The resulting cNMF K-selection plot summarizes model behavior across K values and was used to guide selection of K=4 for downstream analyses. (**B**) GEP activity was summarized across samples using two complementary metrics. Top subpanel shows boxplots of cNMF normalized usage for each sample. In each boxplot, the center line indicates the median usage, boxes denote the interquartile range (25th-75th percentiles), whiskers extend to 1.5* the interquartile range, and points beyond the whiskers represent outliers. Bottom subpanel shows bar plots of fraction of cells assigned to each GEP for each sample. (**C**) Heatmap of pairwise Spearman correlation matrix calculated from GEP signature scores. (**D**) Hallmark gene set enrichment was computed for each cNMF-derived gene expression program (GEP1-GEP4) by performing GSEA on GEP signature scores for each program. Significant pathways (p < 0.05) were retained and visualized as a dot plot with rows representing pathways and columns representing GEPs. Dot color indicates normalized enrichment score (NES), and dot size represents statistical significance as -log10 of Benjamini-Hochberg corrected p values. Pathways were hierarchically clustered based on Pearson correlation distance using average linkage, and the resulting dendrogram is displayed adjacent to the dot plot to highlight pathway modules enriched in shared or distinct GEPs. (**E**) GSEA was performed on Gene Ontology (GO) biological process terms using GEP4 signature scores. The running enrichment score (ES) is shown across the ranked gene list for a set of curated GO terms: negative regulation of gene expression, epigenetic (GO:0045814), regulation of DNA repair (GO:0006282), and mRNA processing (GO:0006397). In each subpanel, the running enrichment score (ES) is plotted across the ranked gene list (top), with tick marks indicating positions of MP genes in the ranked list (middle), and the underlying ranking metric shown below (bottom). The annotated inset reports the normalized enrichment score (NES) and Benjamini-Hochberg corrected p-value for each GO term. (**F**) Heatmap showing expression of cNMF-derived 50 top-scoring genes for each gene expression program (GEP1-GEP4) across Xenium single cells. Normalized Xenium expression values were mean-centered and scaled for visualization. Top annotation indicates binary per-cell GEP assignments. Selected marker genes specific to each GEP are labeled on the right margin for reference.

**Supplementary Figure S7.**
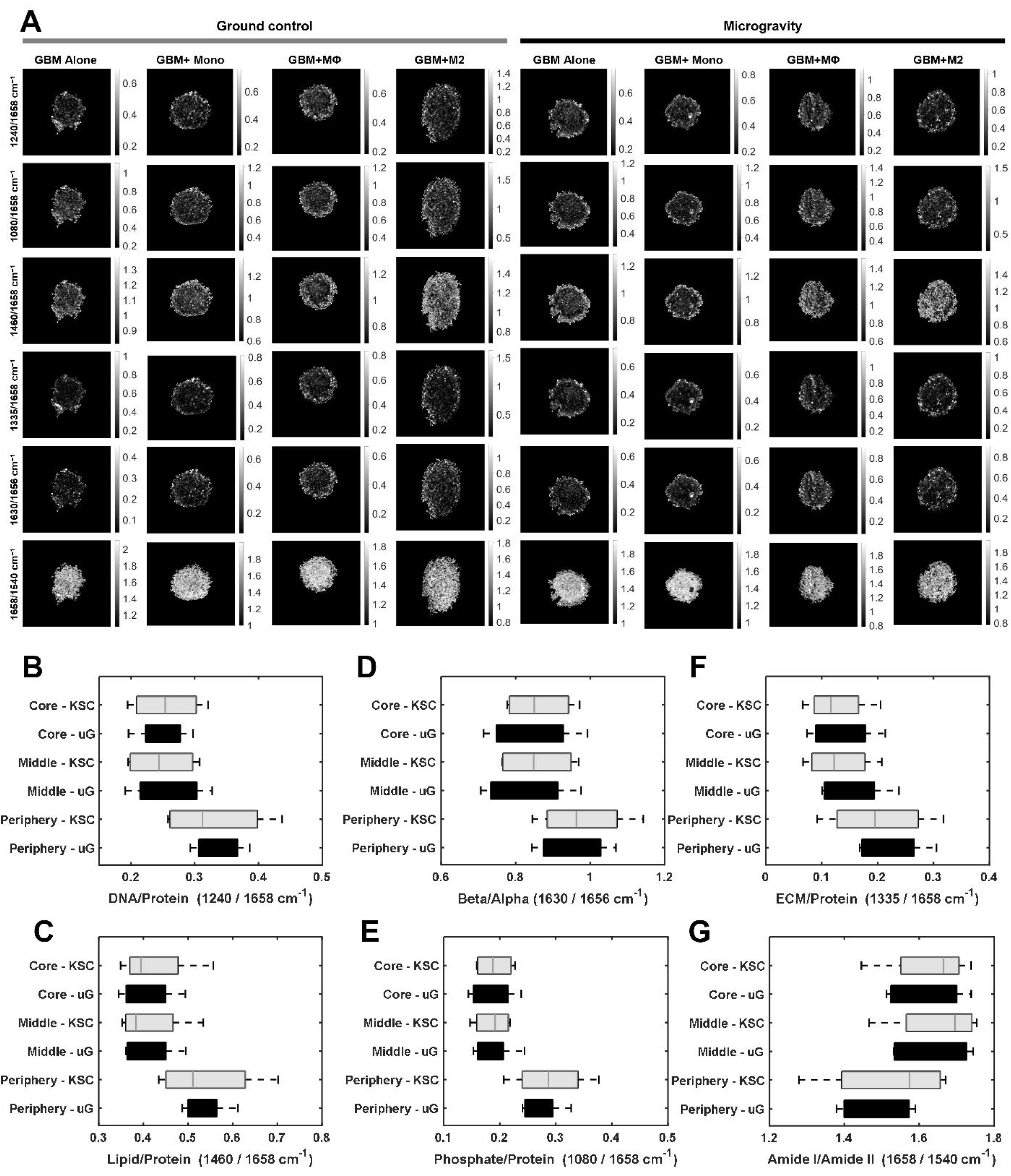
DF-IR images and spatial trends for biologically relevant ratios of spectral bands. (**A**) Band ratio maps of each organoid with intensity values corresponding to the indicated ratio. Box plots of DF-IR band ratios, comparing Ground control and Microgravity across all conditions: (**B**) DNA/Protein (1240/1658 cm⁻¹), (**C**) Lipid/Protein (1460/1658 cm⁻¹), (**D**) Beta/Alpha (1630/1656 cm⁻¹), (**E**) Phosphate/Protein (1080/1658 cm⁻¹), (**F**) ECM/Protein (1335/1658 cm⁻¹), and (**G**) Amide I/ Amide II (1630/1656 cm⁻¹).

**Supplementary Figure S8.**
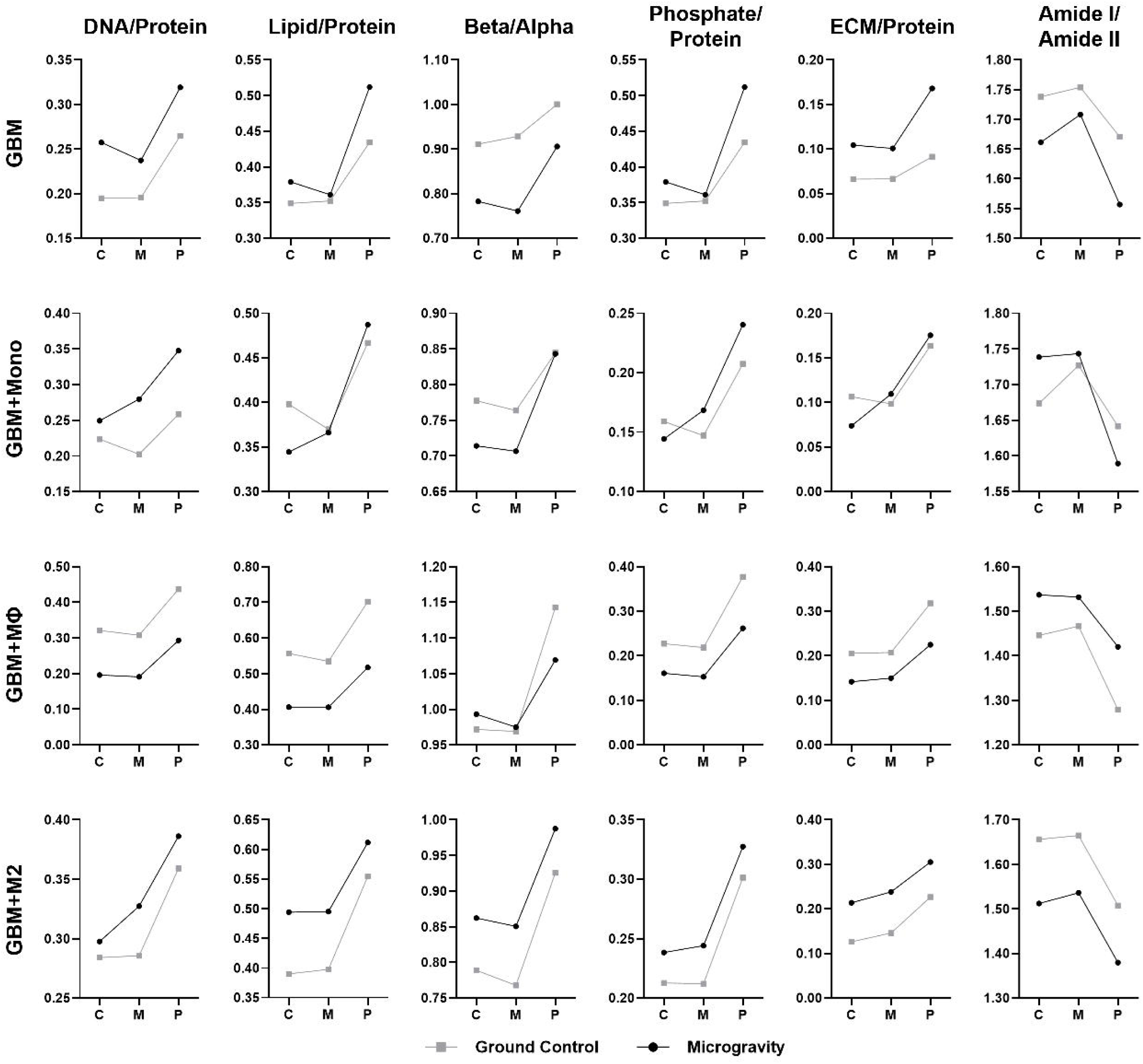
Comparison of radial trends for relevant spectral bands for each organoid condition. The band ratio values for the DNA/Protein, Lipid/Protein, Beta/Alpha proxy, Phosphate/Protein, ECM/Protein, and Amide I/Amide II bands are plotted for each organoid condition, across the core (C), middle (M), and periphery (P) regions.

